# The L27 Domain of MPP7 enhances TAZ-YY1 Cooperation to Renew Muscle Stem Cells

**DOI:** 10.1101/2023.11.01.565166

**Authors:** Anwen Shao, Joseph L. Kissil, Chen-Ming Fan

## Abstract

Stem cells regenerate differentiated cells to maintain and repair tissues and organs. They also replenish themselves, i.e. self-renewal, for the regenerative process to last a lifetime. How stem cells renew is of critical biological and medical significance. Here we use the skeletal muscle stem cell (MuSC) to study this process. Using a combination of genetic, molecular, and biochemical approaches, we show that MPP7, AMOT, and TAZ/YAP form a complex that activates a common set of target genes. Among these targets, *Carm1* can direct MuSC renewal. In the absence of MPP7, TAZ can support regenerative progenitors and activate *Carm1* expression, but not to a level needed for self-renewal. Facilitated by the actin polymerization-responsive AMOT, TAZ recruits the L27 domain of MPP7 to up-regulate *Carm1* to the level necessary to drive MuSC renewal. The promoter of *Carm1*, and those of other common downstream genes, also contain binding site(s) for YY1. We further demonstrate that the L27 domain of MPP7 enhances the interaction between TAZ and YY1 to activate *Carm1*. Our results define a renewal transcriptional program embedded within the progenitor program, by selectively up-regulating key gene(s) within the latter, through the combination of protein interactions and in a manner dependent on the promoter context.

## INTRODUCTION

Stem cells are critical for tissue homeostasis. Depending on the tissue, stem cells either cycle constantly, periodically, or rarely under normal physiological conditions (reviewed in Fuchs and Blau, 2020). Their activities are regulated by their microenvironment or niche. Upon injury, stem cells can enter a faster or longer proliferative state to produce differentiated cells for repair. Like any given cell, stem cells interpret mechanical and biochemical signals to enter or exit the cell cycle, but, in addition, they are tasked with giving rise to differentiated cells and to replenishing themselves, i.e. self-renewal. Here we focus on skeletal muscle stem cells (MuSCs), which are profoundly important to muscle homeostasis and regeneration, as well as to muscle diseases, cancers, and aging (reviewed in Relaix et al., 2021; Sousa-Victor et al., 2022).

The main source of MuSCs is the muscle resident PAX7-expressing (PAX7^+^) cells, also known as satellite cells (Mauro, 1961), as elucidated by lineage tracing (Lepper et al., 2009) and cell ablation studies (Lepper et al., 2011; Murphy et al., 2011; Sambasivan et al., 2011). They are attached to the muscle fiber via the apical adherens junction (AJ) and situated on the basal extracellular matrix (ECM) surrounding the muscle fiber. Loss of the AJ proteins M- and N-cadherins leads to MuSC activation and incorporation into myofiber, as well as increased MuSC numbers (Goel et al., 2017). Loss of the ECM-receptor β1-integrin leads to minimal incorporation of MuSC into the myofiber and loss of MuSCs (Rozo et al., 2016). As cadherins and integrins organize actin, the role of actin in MuSC activity has been investigated. Live imaging of quiescent MuSCs revealed elaborate cellular projections that retract during early activation by injury or myofiber isolation (Kann et al., 2022; Ma et al., 2022). Rac and Rho, two small GTPases regulating actin polymerization at cell projections and cortex, are implicated in regulating MuSC quiescence and activation, respectively (Kann et al., 2022). Moreover, Rho stimulates the nuclear entry of the G-actin sensing co-activator, myocardin-related transcription factor (MRTF), in activated MuSCs. The actin-tethered mechanosensitive Ca^2+^ channel Piezo1, which regulates Rho and MuSC cell projections, also facilitates MuSC activation (Hirano et al., 2023; Ma et al., 2022).

Non-canonical Wnt4 signaling has also been implicated in MuSC quiescence by suppressing the mechano-responsive yes-associated transcriptional co-factor (YAP; Eliazer et al., 2019). YAP and related TAZ (also known as WWTR1, and collectively, YAP/TAZ) are co-activators for the TEAD family (TEAD1-4) of DNA-binding transcription factors that drive cell proliferation (reviewed in Ma et al., 2019; Pan, 2022). YAP overexpression promotes the proliferation of activated MuSCs and myoblasts (Judson et al., 2012; Tremblay et al., 2014). Conditional inactivation of *Yap* in mouse MuSCs (*Yap* cKO) compromises muscle regeneration, whereas *Taz* germline mutant mice do not appear to display regeneration defects (Sun et al., 2017). On the other hand, inactivating *Yap* and *Taz* in MuSCs after muscle injury promotes MuSC quiescence (Silver et al., 2021). By contrast, inactivating *Yap* and *Taz* in MuSCs prior to muscle injury has not been reported. YAP/TAZ are well-recognized mechano-responsive co-activators (reviewed in Panciera et al., 2017), but the players that promote their mechano-sensitivity are incompletely understood in the MuSC.

An evolutionarily conserved regulatory pathway that restrains YAP/TAZ is the Hippo kinase cascade (reviewed in Ma et al., 2019; Pan, 2022). When the Hippo pathway is activated (typically at cell-cell junctions), the most downstream kinases LATS1/2 phosphorylate YAP/TAZ to promote retention in the cytoplasm and proteasomal degradation. In mammals, Angiomotin (AMOT) family members (AMOT, AMOTL1, and AMOTL2, collectively AMOTs) interact with multiple Hippo pathway components (reviewed in Moleirinho et al., 2014). At tight junctions (TJs), they interact with Merlin, an upstream kinase of the Hippo pathway. AMOT can be phosphorylated at serine 175 (S175) by LATS1/2, and lose its actin-binding property. Reciprocally, AMOT can stimulate the kinase activity of LATS1/2. Lastly, AMOTs contain LPTY and PPxY motifs that bind to YAP/TAZ. At all these intersection points, AMOTs act to retain YAP/TAZ at cell junctions, on actin, or in the cytoplasm, and promote their degradation. In certain cell types however AMOT increases the level of nuclear YAP (Yi et al., 2013), a role which appears to be carried out by the non-phosphorylated AMOT (Moleirinho et al., 2017). Whether the non-phosphorylated AMOT also modulates YAP’s transcriptional activity and helps target gene selection is unknown.

Before being identified as associated with the Hippo pathway, AMOT was identified as an angiostatin-binding protein involved in endothelial cell migration and proliferation (reviewed in Moleirinho et al., 2014). In the case of migration, AMOT helps localize Rho activity to the leading edge of endothelial cells. At TJs, AMOT binds and promotes Rich1 (a GTPase activating protein)-mediated hydrolysis of Rac1 and Cdc42 to compromise TJs. Several key TJ proteins, including membrane palmitoylated protein 5 (MPP5, also known as Pals1), interact with AMOT. MPP7, related to MPP5, also exists in an AMOT-containing protein complex (Wells et al., 2006). MPP7 has been implicated in the maintenance of TJs and AJs through binding to MPP5/Crumb (Stucke et al., 2007) and DLG/LIN7 (Bohl et al., 2007), respectively, but AMOT was not included in those studies. On the other hand, we have suggested an unconventional mode of action for MPP7 in MuSC renewal: transcriptional regulation through interaction with AMOT and YAP (Li and Fan, 2017). The mechanisms underlying their tri-partite interaction and target gene activation in MuSCs are not clear.

Although YAP is a well-recognized transcription co-activator, it can also function as a transcriptional repressor of cell cycle inhibitor genes in human Schwann cells (Hoxha et al., 2020). In this context, YAP was localized to genomic regions occupied by the transcription repressor Ying-Yang 1 (YY1) and the enhancer of zeste homolog 2 (EZH2) in the polycomb repressive complex 2 (PRC2). In C2C12 myoblasts, YY1 was shown to inhibit muscle differentiation by repressing myogenic loci (Lu et al., 2013; Wang et al., 2007). During embryogenesis, the inactivation of *Ezh2* in the myogenic lineage led to the de-repression of non-muscle genes (Caretti et al., 2004). A genetic study of *Yy1* function in MuSC revealed that it represses mitochondrial genes during MuSC activation (Chen et al., 2019). Whether and how YAP1 and YY1 coordinate to repress select genes in the MuSC is yet to be explored.

Here we provide evidence that a complex containing MPP7, AMOT, TAZ(YAP), and YY1, activates high levels of *Carm1* expression, which is necessary for MuSC renewal (Kawabe et al., 2012). We determined the roles of *Mpp7*, *Amot*, and *Yap/Taz* in MuSCs for muscle regeneration genetically, and defined their downstream target genes. Promoters of these downstream genes, including that of *Carm1*, contain TEAD binding sites and surprisingly, YY1 binding sites. We dissected the biochemical interactions of the aforementioned proteins and deciphered the mechanism underlying their convergence to *Carm1* activation. Central to the enhanced transcriptional activity of this complex are the AJ-targeting L27 domain of MPP7 and the F-actin-regulated nuclear shuttling of AMOT. Together, we have uncovered an unexpected layer of YAP/TAZ-regulated MuSC renewal, through incorporating MPP7 and AMOT and cooperating with YY1, to activate *Carm1* to a sufficiently high levels that are necessary for self-renewal.

## RESULTS

### *Mpp7* plays a role in SC self-renewal

To determine the role of *Mpp7* genetically, we generated a mouse model with a conditional *Mpp7^flox^* knockout allele (Figure 1A). The loxP sites flank exon 3, which encodes a part of MPP7’s PDZ domain. Cre-mediated recombination predicts a frameshift with an early stop codon. We combined this allele with a *Pax7*-CreER^T2^ allele (*Pax7^CE^*; Lepper et al., 2009) for tamoxifen (TMX) inducible conditional knockout (cKO), referred to as *Mpp7* cKO (Figure 1B); *Rosa26^YFP^* (Srinivas et al., 2001) was included for marking the recombined cells. The cKO efficiency was ∼ 92% based on immunofluorescence (IF) for MPP7 in control and *Mpp7* cKO MuSCs (Figure S1A, B). Thirty days (d) after TMX (without injury), PAX7^+^ MuSC numbers in the tibialis anterior (TA) muscles were similar between control and *Mpp7* cKO animals (Figure S1C). To assess regeneration, we used the procedure outlined in Figure 1B; 5 additional TMX injections after injury were included to further increase cKO efficiency. At 5 days post-injury (dpi), the *Mpp7* cKO had smaller regenerated myofibers and lower PAX7^+^ MuSC density (Figure 1C-F) compared to the control. At 21 dpi (Figure 1G, H; Figure S1D, E), regenerated myofibers remained smaller and PAX7^+^ SC was density lower in the *Mpp7* cKO than those in the control. Using EdU incorporation *in vivo*, we found fewer proliferated YFP^+^ cells in the *Mpp7* cKO than those in the control (Figure 1I; Figure S1F). YFP^+^ MuSCs (isolated by fluorescence-activated cell sorter, FACS, and cultured *in vitro*) also showed a smaller EdU^+^ fraction of the *Mpp7* cKO compared to that of the control, but no difference in programmed cell death (PCD) was found (Figure S1G-I). Lastly, we used a single myofiber culture assay to assess self-renewal, which is based on the relative fractions of three MuSC-derived cell fates: self-renewal (PAX7^+^), progenitor (PAX7^+^MYOD^+^), and differentiation-committed (MYOD^+^) (Figure 1J; Figure S1J). *Mpp7* cKO had fewer progenitor and renewed cells and more differentiation-committed cells, compared to the control. Together, *Mpp7* functions to support MuSC proliferation and self-renewal after injury/activation.

**Figure 1.**
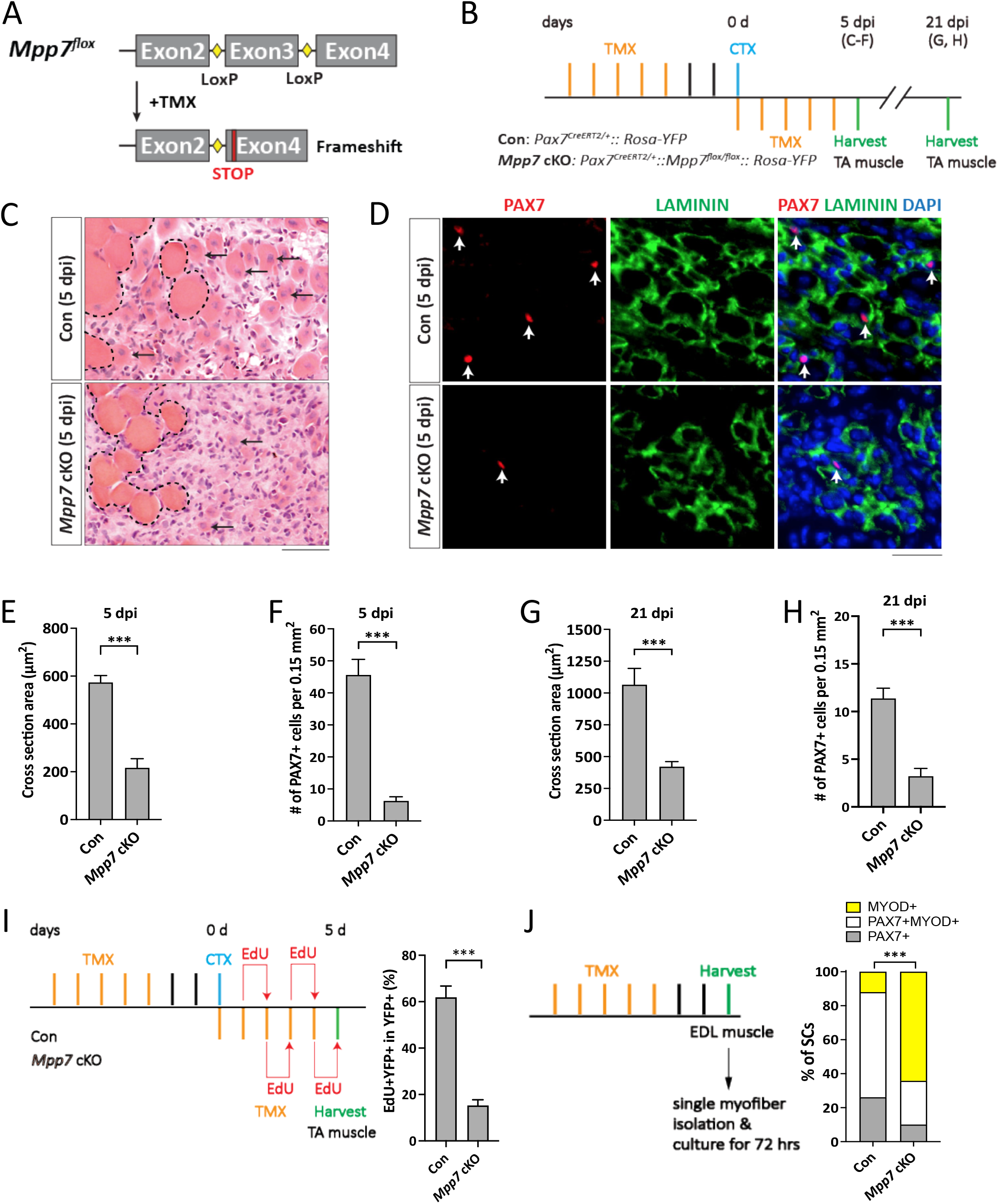
*Mpp7* cKO in Pax7^+^ MuSCs shows defects in regeneration and MuSC self-renewal. (A) Diagram of *Mpp7* floxed allele (*Mpp7^flox^*; loxP, yellow diamond) for tamoxifen (TMX) inducible Cre-mediated cKO. After recombination, out of-frame (Frameshift) joining of Exons 2 and 4 introduces an early stop codon (STOP). (B) Regimen of TMX administration, cardiotoxin (CTX) injury, and tibialis anterior (TA) muscle harvest; d, day; dpi, days post injury. Genotypes of control (Con) and *Mpp7* cKO are indicated. (C-F) *Mpp7* cKO regeneration defects at 5 dpi. Representative images of H&E histology are in (C), immunofluorescence (IF) for PAX7 and LAMININ in (D, with DAPI), and quantification of regenerated myofiber cross sectional area in (E) and of PAX7^+^ MuSC density in (F). Black arrows indicate regenerated myofibers; dashed lines, boundary of injury; white arrows, PAX7^+^ MuSCs. N = 5 mice, each. (G, H) Quantifications of regenerated myofiber cross sectional area (G) and PAX7^+^ MuSC density (H) at 21 dpi. N = 5 mice, each. (I) Regimen of in vivo EdU incorporation to assess the percentage of proliferated YFP-marked cells at 5 dpi; quantification to the right. N = 5 mice, each. (J) Regimen of cell fate determination using single myofiber culture. Cell fates were assessed by IF of PAX7 and MYOD; quantification to the right; keys to cell fates at the top. N = 3 Con mice, of total 601 cells; N = 4 *Mpp7* cKO mice, of total 517 cells. Data information: Scale bars = 25 μm in (C-D). (E-I) Error bars represent means ± SD; Student’s *t*-test (two-sided). (J) Chi-square test. ***, *P*<0.001.

### MPP7’s PDZ and L27 domains are critical for its function in SCs

MPP7 is known to be required for maintaining AJs and TJs in epithelial cells (Bohl et al., 2007; Stucke et al., 2007). We therefore examined whether *Mpp7* inactivation affected apical proteins. Immediately after single myofiber isolation, *Mpp7* cKO MuSCs showed normal levels of apically localized M-cadherin, N-cadherin, β-catenin, and PAR3 (Figure S2A). This is consistent with normal MuSC numbers at 30d after *Mpp7* inactivation.

We next investigated the domains of MPP7 required in the MuSC. MPP7 is composed of an L27, a PDZ, an SH3, and a GUK (last two combined as SH3GUK) domain (Figure 2A; reviewed Chytla et al., 2020). The L27 domain interacts with DLG and LIN7 for AJ targeting (Bohl et al., 2007), the SH3 domain interacts with MPP5 and Crumb for TJ targeting (Stucke et al., 2007), but the PDZ domain has no assigned partner nor known function to date. We transfected expression constructs for full-length (WT), L27-deleted (ΔL27), PDZ-deleted (ΔPDZ), and SH3GUK-deleted (ΔΔSH3GUK) Mpp7 (Figure 2A) into the *Mpp7* cKO MuSCs and assessed their ability to rescue defects in single myofiber culture (Figure 2B). All deletion forms of Mpp7 exhibit similar cellular distribution as WT Mpp7: i.e. nuclear localization in the majority of MuSCs (Figure S2B). WT and ΔΔSH3GUK Mpp7 rescued both progenitor and renewed cells (compared to the control), ΔL27 MPP7 rescued the progenitor but not renewed cells and ΔPDZ MPP7 rescued neither (Figure 2C). Thus, MPP7’s PDZ domain is critical for all protein functions assessed, while the L27 domain is uniquely required for renewal. The SH3 and GUK domains appear to be dispensable.

**Figure 2.**
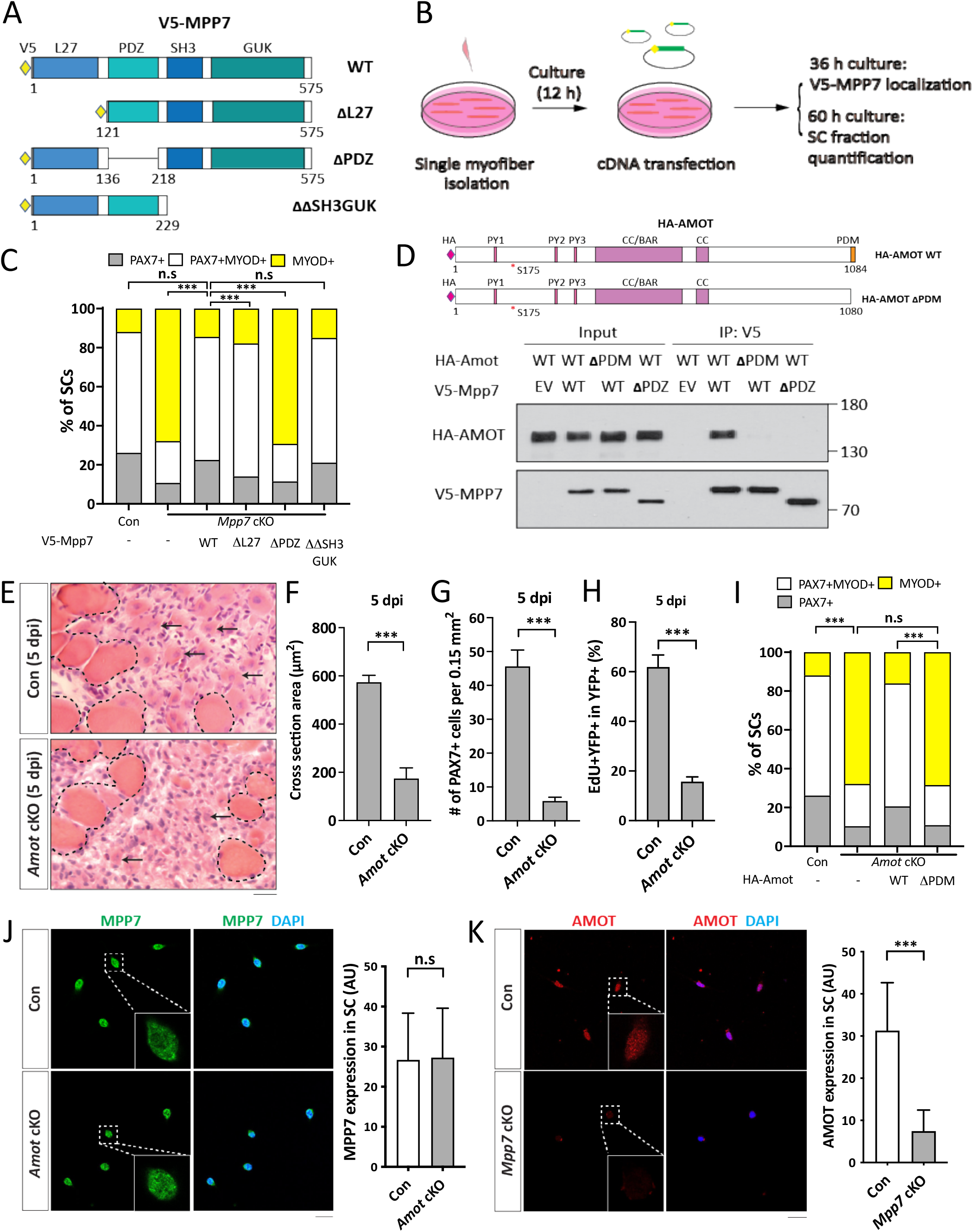
Interacting domains of MPP7 and AMOT are critical for SC renewal. (A) Depiction of MPP7 domain architecture and V5-tagged wild type (WT) and domain deletion mutants (listed on the right) of Mpp7 expression constructs used in (B, C). (B) Flowchart to force-express various Mpp7 constructs (A) in *Mpp7* cKO MuSCs. (C) Quantifications of cells fate from experiments depicted in (B); IF of V5, PAX7 and MYOD was performed to determine the fate of transfected cells. Expression constructs used are in x-axis; (-), empty vector; keys to cell fate at the top. ≥ 168 transfected cells were assessed per group. (D) Co-IP assays to determine interaction domains between MPP7 and AMOT in 293T cells. HA-tagged WT Amot (HA-Amot) and 1′PDM Amot are depicted. V5-tagged WT MPP7 and 1′PDZ MPP7 were used for co-IP with HA-AMOT and 1′PDM AMOT using an anti-V5 antibody, followed by Western blotting with anti-HA or anti-V5 antibodies. (E-H) *Amot* cKO regenerative defects at 5 dpi. Representative H&E staining images of TA muscles from Con and *Amot* cKO are in (E), quantifications of regenerated muscle fiber cross sectional areas in (F), PAX7^+^ MuSC densities in (G), and percentages of EdU^+^ YFP-marked cells in (H). Experimental design is the same as in Figure 1. N = 5 mice, each. (I) Single myofiber transfection assays with WT or △PDM HA-Amot constructs as depicted in (B); (-), empty vector. IF of PAX7, MYOD and HA was performed to determine the fate of transfected cells; keys at the top; ≥ 180 cells per group. (J, K) IF of MPP7 in Con and *Amot* cKO MuSCs in (J) and of AMOT in Con and *Mpp7* cKO MuSCs in (K), at 48 h of culture after FACS isolation. Qualified fluorescent signals (arbitrary units, AU) are to the right of corresponding representative images; 200 cells per group. Data information: Scale bars = 25 µm in (E, J-K). (F-H, J, K): Error bars represent means ± SD. Student’s *t*-test (two-sided). (C, I): Chi-square tests were performed. n.s, *P* > 0.05; ***, *P* < 0.001.

### The PDZ-binding motif (PDM) of AMOT binds to the PDZ of MPP7 and is critical for function

We previously showed an interaction between MPP7 and AMOT by co-immunoprecipitation (co-IP) in 293T cells (Li and Fan, 2017). How they interact and whether *Amot* plays a role *in vivo* were unknown. Using the co-IP assay, we found that MPP7 and AMOT bind to each other via their respective PDZ and PDM domains (Figure 2D; Figure S2E; domain organization of AMOT reviewed in Moleirinho et al., 2014). Importantly, *Amot* cKO mice (same strategy as for *Mpp7* cKO mice described above) showed muscle regeneration defects similar to those of *Mpp7* cKO mice: Smaller myofibers, fewer PAX7^+^ SCs, and reduced EdU incorporation of lineage-marked YFP^+^ cells (Figure 2E-H). In the single myofiber culture, *Amot cKO* MuSCs also had reduced progenitor and renewed cell fractions, which could be rescued by expressing WT Amot but not by ΔPDM Amot (Figure 2I). Because of their interaction, we examined whether their protein levels depended on each other. MPP7 level was not affected in cultured *Amot* cKO MuSCs (Figure 2J). By contrast, the AMOT level was reduced in *Mpp7* cKO MuSCs in culture (Figure 2K) as well as on single myofiber (Figure S2C). Thus, not only the interaction domains of MPP7 and AMOT share a common role in progenitor and renewal fates, their interaction also appears critical to maintain AMOT level (summarized in Figure S2D).

### *Mpp7* cKO and *Amot* cKO MuSCs share differentially expressed genes

We next performed RNA-seq to identify differentially expressed genes (DEGs) in *Mpp7* and *Amot* cKO MuSCs, compared to the control. The experimental design is outlined in Figure 3A and the principle component analysis (PCA) of RNA-seq data is in Figure 3B. *Mpp7* cKO had 58 DEGs and *Amot* cKO had 66 DEGs compared to WT cells (Figure 3C; Table S1). Thirty-five of their DEGs intersect, with 15 of these being downregulated (Figure 3D; Table S1). *Amot* is not a DEG in the *Mpp7* cKO, indicating that its reduced protein levels in this background are post-transcriptionally regulated. As such, some DEGs of *Mpp7* cKO might be a consequence of reduced AMOT levels. GO-term analysis revealed enrichment for estrogen receptor (ESR) signaling, mitochondria biogenesis, and small GTPases (Figure 3E). Genes in our GO-term list have not been studied in the MuSC, except for *Carm1* (or *Prmt4*). CARM1 is an arginine methyl transferase that methylates PAX7, which then recruits epigenetic regulators to activate de novo committed satellite myogenic cells (Kawabe et al., 2012). *Carm1* cKO also has reduced regenerative myofiber size and PAX7^+^ MuSC number. We confirmed that CARM1 level was reduced in *Mpp7* cKO and *Amot* cKO MuSCs (Figure 3F, G), per its downregulated mRNA; the same results were seen in the single myofiber culture (Figure S3A).

**Figure 3.**
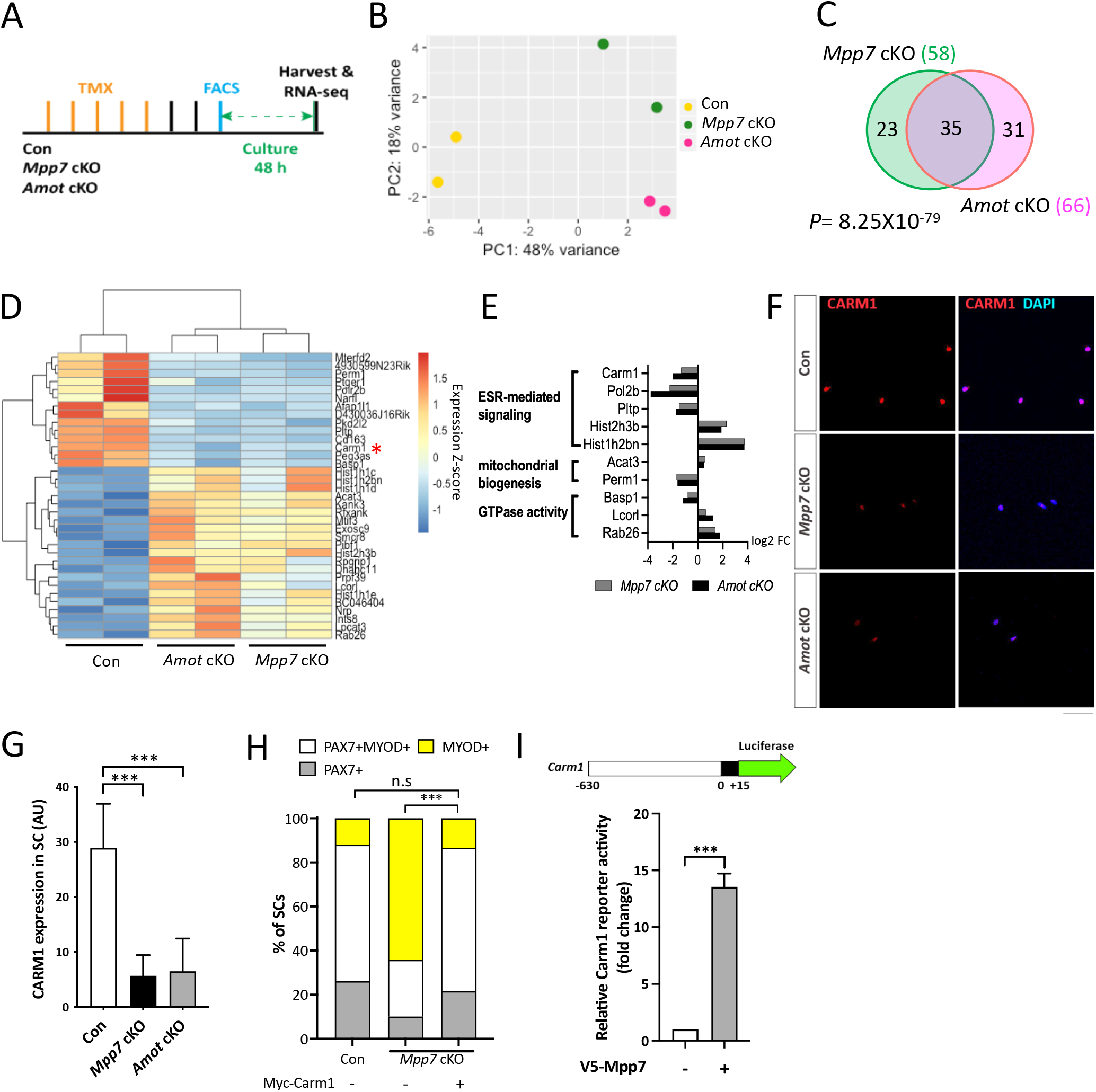
Carm1 is one of the genes commonly regulated by *Mpp7* and *Amot* in the SC. (A) Flowchart for bulk RNA-sequencing (RNA-seq). (B) PCA analysis of transcriptome data of Con, *Mpp7* cKO, *Amot* cKO MuSCs. (C) Venn diagram summarizes (35) overlapping DEGs between the *Mpp7* cKO (58 DEGs) and the *Amot* cKO (66 DEGs). (D) Hierarchical clustering and heatmap of RNA-seq expression z-scores computed for the 35 DEGs in the *Mpp7* cKO and the *Amot* cKO; red asterisk, *Carm1*. (E) Expression fold-changes (log2 FC, log2 fold change) of genes in GO-term enriched pathways (y-axis) are displayed for the *Mpp7* cKO (grey bars) and the *Amot* cKO (black bars). (F, G) Representative IF images of CARM1 in Con, *Mpp7* cKO and *Amot* cKO MuSCs at 48 h in culture are in (F). Quantified CARM1 signals (in AU) are in (G); 200 MuSCs from 2-3 mice in each group. (H) Expressing a Myc-tagged Carm1 (Myc-Carm1) rescues *Mpp7* cKO MuSCs in single myofiber culture; (-), empty vector. IF of Myc, PAX7, and MYOD was performed to determine the fate of transfected cells. Quantification of cell fate fractions are shown; keys at top; ≥ 530 cells in each group. (I) V5-Mpp7 expression construct activates the Carm1-reporter (a luciferase reporter driven by a promoter region (-630 to +15) of *Carm1*, depicted at the top) in 293T cells; (-), empty vector. N=3. Data information: RNA seq data have been deposited in NCBI. Programs used to generate (B, D, E) are described in Methods. Scale bar = 50 μm in (F). Hypergeometric test was used in (C). Error bars represent means ± SD. One-way ANOVA with Tukey’s post hoc test was performed in (G), Chi-square test in (H), and Student’s *t*-test (two-sided) in (I). n.s, *P* > 0.05; ***, *P* < 0.001.

To determine if *Carm1* plays a role in the *Mpp7-*operated pathway, we force-expressed Carm1 in *Mpp7* cKO MuSCs in single myofiber culture and found that it was sufficient to rescue progenitor and renewed fates (Figure 3H). We next tested whether *Mpp7* could regulate *Carm1* transcription. For this, we made a luciferase reporter fused to a putative promoter region (-630 to +15 bp) of *Carm1* (i.e., a Carm1-reporter) and found that overexpression of Mpp7 could activate the Carm1-reporter in 293T cells (Figure 3I). By contrast, Mpp5 could not activate the Carm1-reporter, indicating a selectivity for Mpp7 (Figure S3B). In addition, ΔΔSH3GUK Mpp7 activated the reporter similarly to the WT Mpp7, ΔL27 Mpp7 weakly activated, and ΔPDZ Mpp7 did not activate the reporter (Figure S3C). Thus, among the DEGs of *Mpp7* cKO, *Carm1* is the chief effector gene for progenitor and renewal fates.

### The regulatory network of *Yap* and *Taz* overlaps with those of *Mpp7* and *Amot*

*Mpp7* knock-down reduced nuclear YAP in myoblasts, suggesting MPP7 increases YAP activity (Li and Fan, 2017). AMOT is known to bind to YAP/TAZ directly, and can either increase or decrease nuclear YAP/TAZ in different contexts (reviewed in Moleirinho et al., 2014). Following these threads, we examined an existing set of DEGs obtained by overexpressing YAP or TAZ in myoblasts (Sun et al., 2017)(Figure S3D). We found very few of those overlap with DEGs of *Mpp7* cKO and/or *Amot* cKO, likely due to different experimental approaches. We next asked whether DEGs of *Mpp7* or *Amot* cKOs harbor TEAD-binding sites in their promoters. Indeed, most of their promoters have TEAD-binding sites (Fishilevich et al., 2017; Keenan et al., 2019), including that of *Carm1* (Figure S3E). This encouraged us to employ *Taz^flox^* and *Yap^flox^* (Reginensi et al., 2013) as double cKO in the MuSC (*YapTaz* cKO) and compare with *Mpp7* cKO and *Amot* cKO.

We confirmed that *Yap* cKO had compromised muscle regeneration (Sun et al., 2017), and found that *YapTaz* cKO had a more severe defect (Figure 4A). Previously, *Taz* mutant was reported to have no muscle regeneration defects (Sun et al., 2017), and our *YapTaz* cKO data indicate that *Taz* partially compensates for *Yap* in muscle regeneration. To not miss DEGs due to compensation, we proceeded with RNA-seq using *YapTaz* cKO MuSCs. The *YapTaz* cKO showed 564 DEGs, compared to the control (Figure 4B). As expected, many cell growth- and proliferation-associated pathways were impacted (Figure S4A). Thirty-three *Mpp7* cKO DEGs and 31 *Amot* cKO DEGs overlapped with *YapTaz* cKO DEGs, and 19 were common in all 3 sets of DEGs (Figure 4B; Figure S4B). During our search for transcription factor binding sites, we noticed a congruence of TEAD- and YY1-binding sites in these DEGs’ promoters (Figure S4C). These analyses suggest that MPP7 and AMOT help YAP and/or TAZ to selectively regulate a small set of downstream genes, and YY1 is likely involved in their regulation (see below).

**Figure 4.**
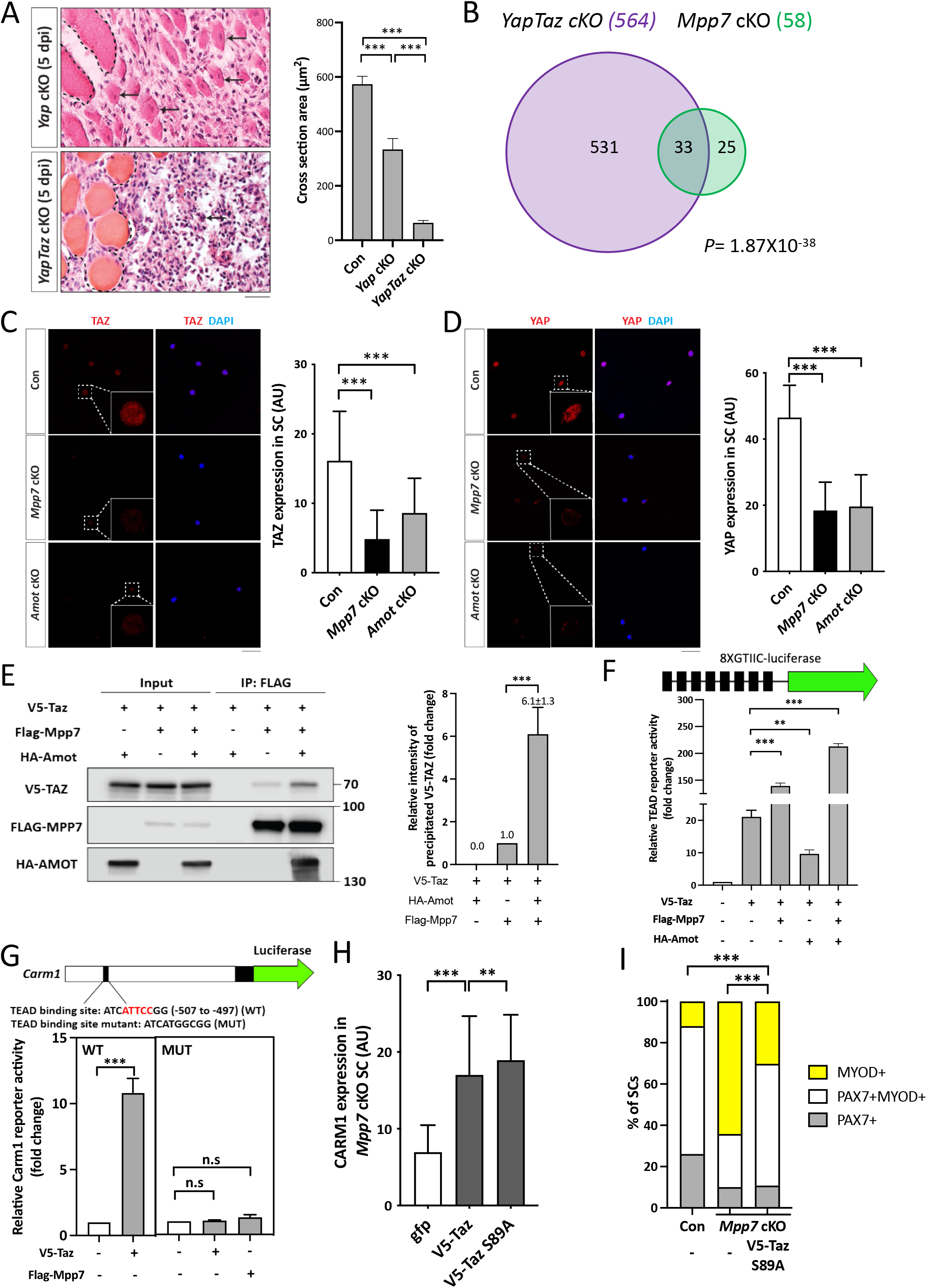
*Mpp7/Amot* regulatory network intersects with that of *Yap/Taz*. (A) Representative H&E histology of *Yap* cKO and *YapTaz* cKO muscles at 5 dpi (Con histology not included); quantifications of regenerated myofiber cross sectional area to the right. N = 5, each genotype. (B) Venn diagram shows overlapping DEGs between the *YapTaz* cKO and the *Mpp7* cKO. (C, D) Representative IF images of TAZ (C) and YAP (D) in FACS-isolated and cultured Con, *Mpp7* cKO, and *Amot* cKO MuSCs at 48 h; quantified fluorescent signals (AU) to the right of corresponding images; 200 MuSCs from 2-3 mice in each group. (E) Co-IP of V5-TAZ and HA-AMOT by FLAG-MPP7 expressed in 293T cells. Expression constructs and tagged epitopes for detection are indicated; (-), empty vector. Quantification of relative levels of co-IPed V5-TAZ is to the right. N = 3. (F) Relative TEAD-reporter (8XGTIIC-luciferase, depicted at top) activities when co-transfected with V5-Taz, Flag-Mpp7, and/or Ha-Amot expression constructs in 293T cells; (-), empty vector. N =3. (G) Relative activities of WT and TEAD-binding site mutated (MUT) Carm1-reporters co-transfected with V5-Taz or Flag-Mpp7 expression constructs; (-), empty vector. N = 3. (H) Quantified IF signals (AU) of CARM1 in *Mpp7* cKO MuSCs transfected with gfp (as control), V5-Taz WT and V5-Taz S89A expression constructs; 200 MuSCs from 2-3 mice in each group. (I) Comparison of cell fate fractions among Con, *Mpp7* cKO, and *Mpp7* cKO MuSCs transfected with V5-Taz S89A expression construct in single myofiber culture; (-), empty vector; keys at the top; ≥ 217 cells in each group. Data information: Scale bar = 25 µm in (A) and 50 µm in (C, D). Error bars represent means ± SD. Hypergeometric test was used in (B); One-way ANOVA with Tukey’s post hoc test was performed in (A, C, D, F-H); Chi-square test in (I). n.s, *P* > 0.05; **, *P* < 0.01; ***, *P* < 0.001.

Neither *Yap* nor *Taz* were a DEG in *Mpp7* or *Amot* cKOs, but their protein levels were reduced in both cKOs; with TAZ levels being the most reduced in *Mpp7* cKO MuSCs (Figure 4C, D). The addition of proteasome inhibitor MG132 to *Mpp7* cKO MuSCs restored the levels of TAZ, YAP, and AMOT near to those in the control, but the level of CARM1 was only partially restored (Figure S4D). This is puzzling if *Carm1* is a target of YAP/TAZ (via TEAD). We reasoned that MPP7 must have a role other than stabilizing TAZ, YAP, and AMOT. Using TAZ as a representative for TAZ and YAP (because TAZ level is most affected in *Mpp7* cKO MuSCs), we showed that MPP7 could interact with TAZ, and their interaction was enhanced by AMOT (Figure 4E; Figure S4F). Using a TEAD-reporter (Dupont et al., 2011), we found that MPP7 co-expression substantially increased reporter activation by TAZ, and adding AMOT further boosted the reporter activity (Figure 4F). AMOT alone however reduced TAZ’s activity, consistent with it being a negative regulator in most studies. We conclude that MPP7 reverses AMOT’s negative role for TAZ, and MPP7 and AMOT together facilitate TAZ function as a co-activator.

We next asked if the TEAD binding site in the *Carm1* reporter mediated the responsiveness to Mpp7. The *Carm1*-reporter with the TEAD binding site mutated was no longer responsive to Mpp7 (Figure 4G). Given that Carm1 is sufficient to rescue the defects of *Mpp7* cKO MuSCs, we asked whether *Taz* could also do so. Both TAZ and TAZ S89A (a stabilized form of TAZ; Kanai et al., 2000) could only increase CARM1 levels by ∼2-fold in *Mpp7* cKO MuSCs, and TAZ S89A performed slightly better than TAZ (Figure 4I; Figure S4E). However, TAZ S89A could not rescue the renewal fate of *Mpp7* cKO MuSCs, even though it rescued the progenitor fate (Figure 4J). Thus, while TAZ can up-regulate *Carm1* to rescue the progenitor fate, MPP7 is additionally needed to boost *Carm1* expression to a higher level required for the renewal fate.

### L27 domain of MPP7 enhances TAZ-mediated transcription and SC renewal

To dissect the mechanism underlying MPP7-enhanced TAZ activity, we examined the domains of MPP7 required for their synergy. By co-IP, we found that ΔPDZ MPP7 failed to interact with TAZ, ΔL27 MPP7 had a diminished interaction with TAZ, and ΔΔSH3GUK MPP7 had the same level of interaction with TAZ as WT MPP7 (Figure 5A). When tested on the TEAD-reporter, both ΔPDZ Mpp7 and ΔL27 Mpp7 had little activity, even though a low level of interaction was observed for ΔL27 MPP7 and TAZ (Figure 5B). The interaction between MPP7 and TAZ is likely indirect via endogenous AMOT in 293T cells: 1) ΔPDZ MPP7 cannot bind AMOT; 2) ΔPDM AMOT, not able to bind MPP7, interfered with MPP7-TAZ interaction, likely by competing TAZ away from endogenous AMOT (Figure S5A); 3) TAZ with a mutated WW domain (TAZ WWm) unable to interact with AMOT, failed to interact with MPP7 (Figure S5B).

**Figure 5.**
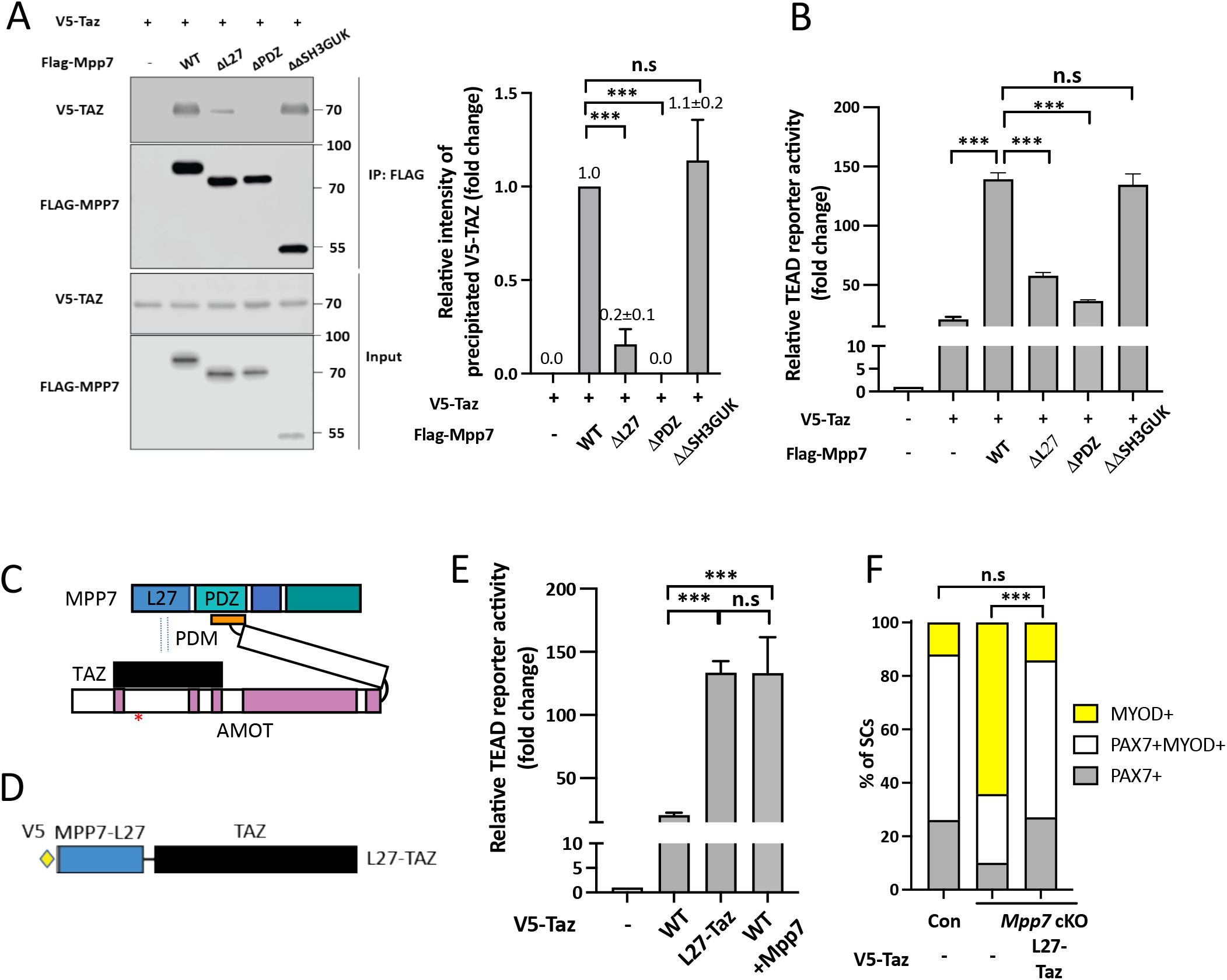
AMOT links TAZ to the MPP7-L27 domain for enhanced transcriptional activity. (A) MPP7-L27 contributes to MPP7-TAZ interaction by co-IP assay in 293T cells. Expression constructs and tagged epitopes for detection are indicated; (-), empty vector. Quantification of co-IPed V5-TAZ are to the right. N = 3. (B) Relative TEAD-reporter activities when co-transfected with various Mpp7 expression constructs (x-axis); (-), empty vector. N = 3. (C) A model summarizes the co-IP results in Figures 5A, S5A and S5B. (D) Diagram of the fusion construct between V5-tagged Mpp7-L27 and Taz (i.e., L27-Taz) used in (E, F). (E) Relative TEAD-reporter activities when co-transfected with expression constructs indicated in the x-axis; (-), empty vector. N = 3. (F) Quantification of cell fates of Con and *Mpp7* cKO MuSCs in single myofiber assay; expression constructs in the x-axis; (-), empty vector; ≥ 155 cells in each group. Data information: Error bars represent means ± SD. One-way ANOVA with Tukey’s post hoc test was performed in (A, B, E), and Chi-Squire test, in (F). n.s, *P* > 0.05; **, *P* < 0.01; ***, *P* < 0.001.

The above results suggest that the L27 domain of MPP7 (MPP7-L27) enhances the interaction between MPP7 and TAZ only when they are brought into proximity via two distinct parts of AMOT (Figure 5C), and the interaction between MPP7-L27 and TAZ is key to enhancing TAZ’s transcriptional activity. If so, a fusion of MPP7-L27 to TAZ (L27-TAZ; Figure 5D) should bypass their interaction via the obligatory AMOT and have enhanced transcriptional activity. Indeed, L27-TAZ activated the TEAD-reporter to the same level as TAZ and MPP7 together (Figure 5E). L27-TAZ also activated the Carm1-reporter effectively (Figure S5C). We further dissected MPP7’s L27 domain, which consists of two L27 repeats, L27N and L27C. L27N-TAZ and L27C-TAZ fusions showed slightly reduced transcriptional activity than the full L27-TAZ. The L27N binds DLG and L27C, to LIN7 (Bohl et al., 2007). Mutations of Lysine 38 (L38) in L27N and L95 in L27C should interrupt interactions with DLG and LIN7, respectively. Yet neither mutation in L27-TAZ affected transcriptional activity (Figure S5C), indicating that DLG and LIN7 are not involved in this context. Importantly, L27-TAZ increased CARM1 levels in MuSCs more than TAZ did (Figure S5D), and rescued the progenitor and renewal fates of *Mpp7* cKO SCs (Figure 5F). Uncovering the role of MPP7-L27 for MuSC renewal by enhancing TAZ’s transcriptional activity brings us two outstanding questions: 1) Is AMOT’s role only to bring TAZ (YAP) to the proximity of MPP7-L27? 2) How does MPP7-L27 enhance TAZ function?

### AMOT acts as a F-actin-regulated shuttling factor in MuSCs

AMOT is proposed to function as a scaffold protein to promote the localization of YAP to the cytoplasm, AJs, TJs, and F-actin, thereby precluding nuclear YAP. However, in some cell types AMOT promotes nuclear YAP (reviewed in Moleirinho et al., 2014). When MuSCs were isolated on single myofibers in the presence of a ROCK inhibitor Y-27632 (Kann et al., 2022), AMOT, MPP7, and YAP (TAZ not detectable) were localized to the apical side and cell projections (Figure 6A); ROCK acts downstream of Rho to promote actin polymerization. We next subjected MuSCs to pharmacological reagents that affect actin polymerization (Figure 6B) and examined the localization of AMOT and MPP7. Immediately after FACS isolation, MuSCs treated with Jasplakinolide (Jasp) and Narciclasine (Nar) to stabilize F-actins had increased nuclear AMOT and MPP7, compared to mock-treated cells (Figure 6C). Conversely, MPP7 and AMOT were mostly nuclear in activated MuSCs at 48 h of culture but shuttled out to the cytoplasm when treated with Blebbistatin (Bleb), Cytochalasin B (Cyto B), or Y-27632 to weaken or disrupt F-actin (Figure 6D; Figure S6A).

**Figure 6.**
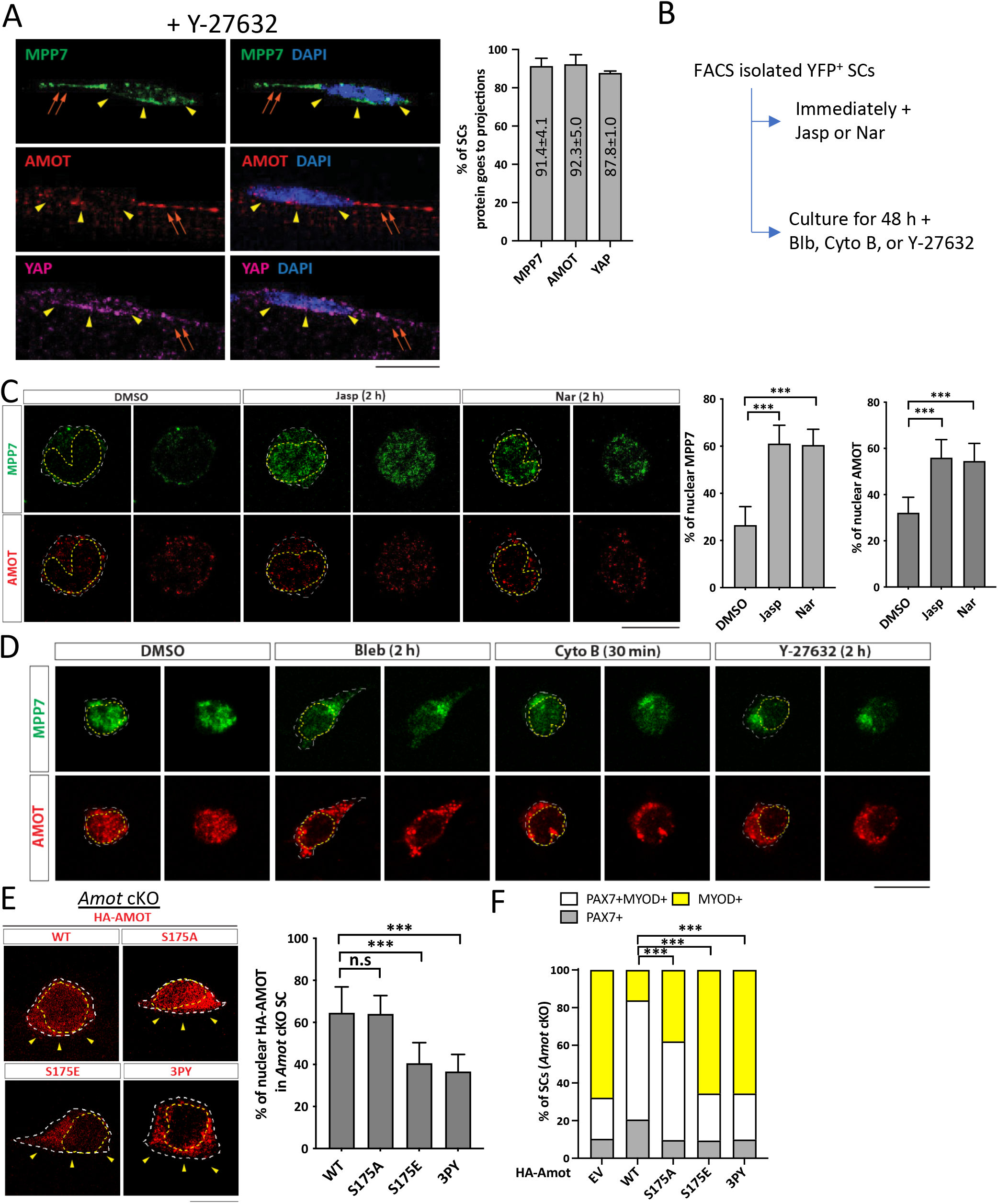
Actin polymerization state impacts AMOT localization and function through binding to TAZ or YAP. (A) Representative IF images of MPP7, AMOT, and YAP in wild type MuSCs on single myofibers isolated in the presence of 50 µM Y-27632. Percentages of MuSCs with MPP, AMOT and YAP signals in cellular projections were quantified: 77 cells for MPP7, 73 cells for AMOT, and 66 cells for YAP, from 3 mice. (B) Experimental flowchart to investigate the impact of actin polymerization state on the cellular localization of MPP7 and AMOT in FACS-isolated MuSCs in culture. (C) Representative IF images of MPP7 and AMOT of freshly isolated MuSCs treated with DMSO (control), 100 nM Jasplankinolide (Jasp), or 100 nM Narciclasine (Nar) for 2 h. Percentages of MPP7 or AMOT IF signals in the nucleus (versus total signals) in each SC were quantified; 200 MuSCs in each group. (D) Representative IF images of MPP7 and AMOT in MuSCs cultured for 48 h and treated with DMSO (control), 10 µM Blebbinstatin (Bleb), 10 µM Cytochalasin B (Cyto B), or 10 µM Y-27632 for indicated time prior to assay. See Figure S6A for quantified data. (E) Localization of HA-tagged AMOT WT, AMOT S175A (S175A), AMOT S175E (S175E), and AMOT 3PY (3PY) expressed (via transfection) in *Amot* cKO MuSCs on single myofibers by IF of HA; yellow arrowheads, apical side. Percentages of nuclear signals (of total signal) of each variant were quantified; 50 MuSCs in each group. (F) HA-tagged AMOT variants in (E) were transfected into *Amot* cKO SCs on single myofibers; (-), empty vector. Cell fates of transfected cells were determined by IF of HA, PAX7 and MYOD and quantified; ≥ 145 cells in each group. Data information: Scale bars = 10 µm in (A, C-E). Error bars represent means ± SD. One-way ANOVA with Tukey’s post hoc test was performed in (A, C, E), and Chi-square test in (F). n.s, *P* > 0.05; ***, *P* < 0.001.

Curiously, MPP7 and AMOT did not show strict co-localization in the MuSC, but their cytoplasmic versus nuclear localization are similarly regulated by actin states and MPP7 helps stabilize AMOT. The AMOT-MPP7 complex is likely dynamic, while their cell compartment localization depends on AMOT and F-actin interaction. AMOT and F-actin interaction (Ernkvist et al., 2006) is modulated by phosphorylation of serine 175 (S175): S175A AMOT can bind actin, whereas phosphor-mimetic S175E AMOT cannot (Chan et al., 2013; Dai et al., 2013). We expressed both forms of AMOT in *Amot* cKO MuSCs. Similar to those reported in cell lines (Moleirinho et al., 2017), higher percentages of WT and S175A AMOT were present in the nucleus compared to S175E AMOT in transfected MuSCs (Figure 6E). Unlike WT Amot, S175A Amot could only rescue the progenitor but not the renewal fate; S175E Amot bore no rescue activity (Figure 6F). Furthermore, AMOT binding to TAZ/YAP is critical for MuSC renewal, as AMOT with all 3 TAZ/YAP binding motifs mutated (denoted as 3PY) was largely cytoplasmic (Figure 6E) and did not rescue *Amot* cKO MuSCs (Figure 6F). These results together suggest that the dynamic association of AMOT with F-actin (and with MPP7), instead of a strictly phosphorylated or non-phosphorylated form of AMOT per se, is critical for MuSC renewal, and its binding to YAP/TAZ is indispensable.

### YY1 adds another dimension to MPP7-L27 and TAZ for transcriptional activity

We next addressed how MPP7-L27 and TAZ acquire higher transcriptional activity than TAZ alone. As mentioned before, YY1 binding sites are prevalent in the promoter regions of genes commonly regulated by MPP7, AMOT, and YAP/TAZ (Figure S4B). The *Yy1* cKO has been shown to cause dysregulated mitochondrial and glycolic genes in MuSCs (Chen et al., 2019). We found significant, though not extensive, overlaps between our DEGs and *Yy1* cKO DEGs (Figure S7A). Four examples of the promoters of these overlapping DEGs are depicted in Figure 7A.

**Figure 7.**
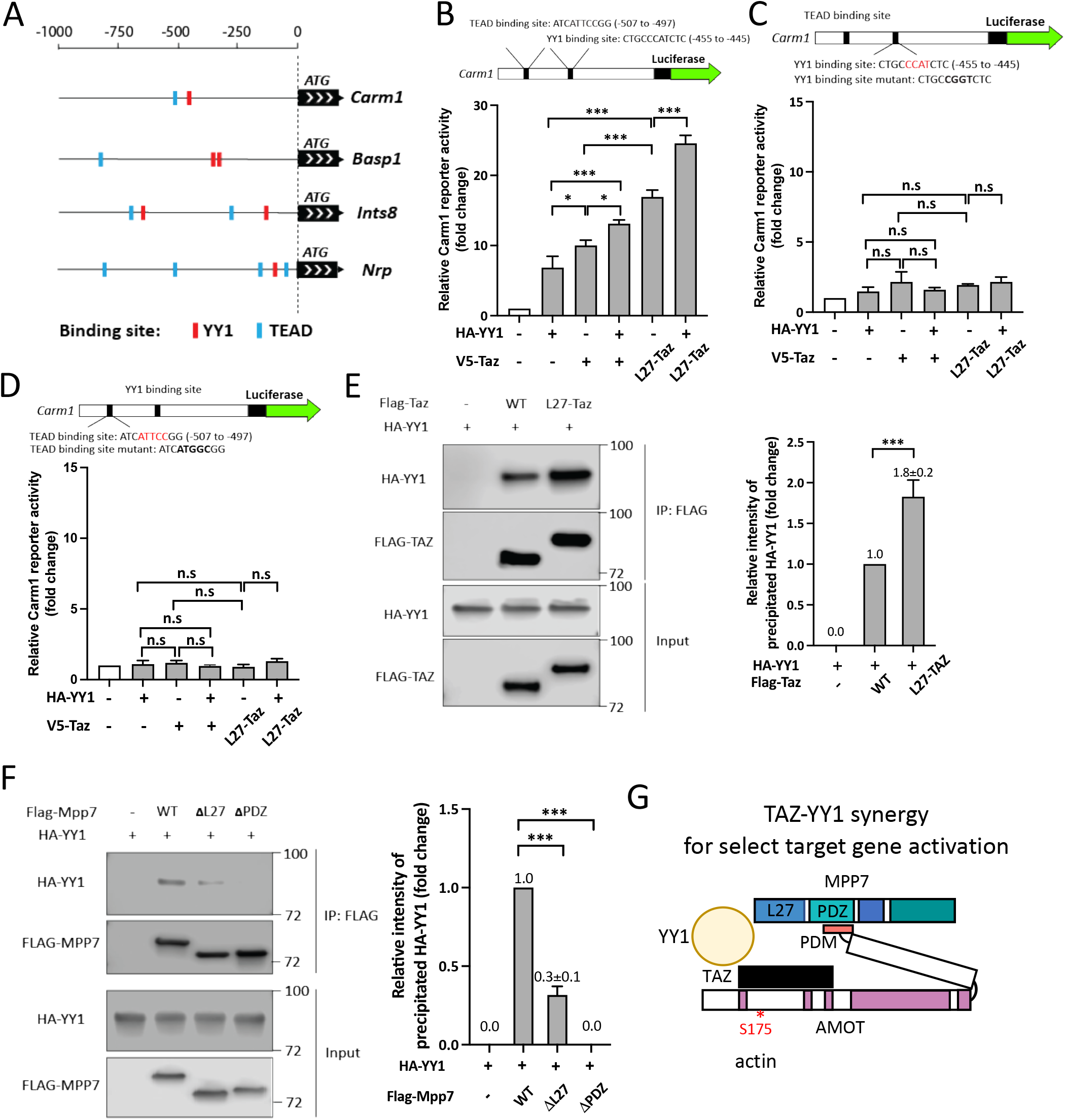
MPP7’s L27 domain cooperates with TAZ and YY1 to enhance transcription. (A) Schematics for promoter regions of 4 DEGs among *Mpp7, Amot, YapTaz,* and *Yy1* cKO (Chen et al., 2019) data sets. Putative YY1 binding sites are shown as red blocks and TEAD binding sites, blue blocks. (B) The location and sequence of the TEAD and YY1 binding sites are indicated in the *Carm1* promoter. Carm1-reporter was co-transfected with expression constructs in x-axis to assess transcriptional activity; (-), empty vector. N = 3. (C) Same as in (B), except that the YY1 binding site was mutated (top). N=3. (D) Same as in (B), except that the TEAD binding site was mutated (top). N=3. (E) Co-IP assay shows the interaction between TAZ and YY1 can be enhanced by fusing MPP7’s L27 domain to TAZ in 293T cells. Expression constructs and tagged epitopes for detection are indicated. Quantification of co-IPed HA-YY1 is to the right. N = 3. (F) Co-IP assay to determine the relative contributions of L27 and PDZ domains of MPP7 to its interaction with HA-YY1 in 293T cells. Expression constructs and tagged protein detection are indicated. Quantification of co-IPed HA-YY1 is shown to the right. N = 3. (G) A model for TAZ-YY1 cooperation mediated by AMOT and MPP7. S175 of AMOT is subjected to phosphorylation and regulation by actin dynamics. Data information: Error bars represent means ± SD. One-way ANOVA with Tukey’s post hoc test was performed in (B-F). n.s, *P* > 0.05; *, *P* < 0.05; ***, *P* < 0.001.

To test the relevance of YY1, we examined its ability to activate the Carm1-reporter. Both TAZ and YY1 can activate the reporter, and L27-TAZ displayed higher activity than TAZ with or without YY1, but no synergy was observed (Figure 7B). When we mutated either the TEAD or the YY1 binding site, no combinations of YY1, TAZ, and L27-TAZ showed any appreciable activity (Figure 7C and D); WT Mpp7 also could not activate the mutated Carm1-reporters (Figure S7B). One explanation is that exogenous YY1 or TAZ activates the promoter by cooperating with endogenous TAZ (or YAP) and YY1, respectively. That is, their ubiquitous expression masks their cooperativity, and their co-existing binding sites enhance the efficiency of co-occupancy to activate transcription.

To have a higher efficiency of promoter co-occupancy, they likely interact with each other. Indeed, TAZ and YY1 could interact with each other by co-IP, and L27-TAZ performed better than TAZ (Figure 7E). Furthermore, YY1 interacted with MPP7 indirectly through endogenous AMOT, as their interaction also required the AMOT-binding PDZ domain of MPP7 (Figure 7F). YY1-MPP7 interaction (through endogenous AMOT) was also diminished when the L27 domain was deleted, consistent with its diminished interaction with TAZ. Although L27-TAZ and YY1 together activated the Carm1-reporter the most, we could not exclude a possible contribution of AMOT in transcriptional activity as TAZ could recruit AMOT. To test this, we made an L27-TAZ WWm (which precludes AMOT binding) and found it to activate the reporter equally well as L27-TAZ, either with or without YY1 (Figure S7C). We summarized these findings in Figure 7G and propose that the MPP7-AMOT-TAZ(YAP)-YY1 protein complex is a stronger activation complex targeting a select set of genes intersecting the YY1 and YAP/TAZ pathways. Although AMOT does not contribute to the transcriptional activation, it is key to bridging the complex together, and its sensitivity to the actin polymerization state endows this complex to respond to mechanical changes of MuSCs from quiescence to activation.

## Discussion

Here we show that an array of protein interactions converges to upregulate *Carm1* to a level necessary for MuSC self-renewal. MPP7, AMOT, and YAP are localized to quiescent MuSCs’ lateral projections, poised for nuclear entry. As the actin cytoskeleton undergoes re-arrangement during activation, AMOT and MPP7 enter the nucleus and help stabilize nuclear YAP/TAZ. The L27 domain of MPP7 plays a central role in enhancing the interaction between TAZ and YY1 on the *Carm1* promoter for high levels of expression. We propose that MuSC renewal through upregulating *Carm1* is tied to a mechano-responsive mechanism in the damaged and regenerative muscle environment where MuSCs experience a dynamic flux of mechanical forces.

### Tiered regulation of *Carm1* levels for MuSC renewal

Here we uncovered a mechanism underlying transcriptional regulation of *Carm1* in MuSC renewal. CARM1 was first described as having the ability to methylate PAX7 thus enabling activation of de novo MuSCs during asymmetric division (Kawabe et al., 2012), and its activity was regulated by p38 MAP kinase (Chang et al., 2018). Our RNA-seq data reveal that *Carm1* is a shared DEG among the cKOs of *Mpp7*, *Amot*, *Yap/Taz*, and *Yy1*. We focused on *Carm1* in this study, but other common DEGs may also contribute to MuSC activity. All the above cKOs, including *Carm1* cKO (Kawabe et al., 2012) display muscle regeneration defects. In particular, *Mpp7* and *Amot* support MuSC proliferation and renewal in vivo and in vitro, as does *Carm1* (Kawabe et al., 2012). *Carm1* promoter harbors indispensable TEAD and YY1 binding sites for transcriptional activation, and MPP7 contributes to Carm1 regulation by providing its L27 domain, via AMOT as an intermediate bridge protein, to facilitate TAZ-YY1 interaction.

Curiously, most *YapTaz* cKO’s DEGs are unaffected in either *Mpp7* cKO or *Amot* cKO. We reason that the remaining levels of YAP/ TAZ in *Mpp7* cKO or *Amot* cKO are sufficient to activate those non-overlapping DEGs to allow a reduced level of muscle regeneration. Such levels of YAP/ TAZ are however insufficient to activate high levels of *Carm1*, leading to reduced progenitors and renewed MuSCs. Without YAP and TAZ, i.e. *YapTaz* cKO, cell growth and proliferation are severely compromised, leading to a severe regenerative deficit. Our results extend previous studies of YAP and/or TAZ (Silver et al., 2021; Sun et al., 2017; Tremblay et al., 2014) concerning the proliferative state of MuSCs, by including MPP7, AMOT, and YY1, and linking them to regulate *Carm1* transcription. While TAZ(YAP) can activate *Carm1*, it needs to cooperate with MPP7 (via AMOT) and YY1 to activate *Carm1* to a higher tier of expression level to drive de novo generation of satellite myogenic cells.

### AMOT is a conduit for YAP and TAZ’s mechano-responsive transcriptional regulation

AMOT and MPP7 bind each other, their cKO MuSCs share many DEGs, and their nuclear versus cytoplasmic localization is co-regulated by actin polymerization states. Given AMOT’s association with F-actin and Rich1, suggests it plays the actin-sensor role within the AMOT-MPP7 partnership. Reciprocally, MPP7 plays a chaperone-like role in the stability of AMOT. Their IF signals are not strictly co-localized in the MuSC, and yet their interaction was visualized by PLA (Li and Fan, 2017). Furthermore, WT Amot can rescue *Amot* cKO MuSC renewal, but ΔPDM-Amot, F-actin-binding-able S175A Amot, F-actin-binding-disabled S175E Amot, and TAZ/YAP-binding-defective 3PY Amot cannot. Together, our results suggest that dynamic association with MPP7 and F-actin, as well as binding to YAP/TAZ are all critical for AMOT function.

Recent work revealed that actin regulation by Rac at the MuSC’s lateral projections and by Rho at the cortex controls MuSC’s quiescence and activation, respectively (Kann et al., 2022). There, the nuclear entry of MRTF, a G-actin sensing co-activator, was found to respond to Rho activation. Nuclear YAP is also regulated by Rho activation (Dupont et al., 2011). Our results showing that AMOT can respond to actin polymerization states and bring MPP7 into the nucleus to do YAP/TAZ’s bidding, support that AMOT is a direct mechano-sensor for YAP/TAZ’s mechano-responsive transcription. The distinct and overlapping transcriptomes governed by YAP/TAZ and MRTF have been explored in cancer cells (Foster et al., 2017). How they operate inter-dependently in the MuSC is of great future interest.

### The L27 domain of MPP7 plays a key role in MuSC renewal

We reported that MPP7 was detected in activated MuSC nuclei (Li and Fan, 2017), contrasting prior reports of its localization at AJ and TJ (Bohl et al., 2007; Stucke et al., 2007). Our data here indicate that multiple regions of MPP7 can mediate its nuclear entry, possibly via other partner proteins. While the essential interaction between MPP7-PDZ and AMOT-PDM motif is predictable, the role of its L27 domain for MuSC renewal and enhancing TAZ’s transcriptional activity is unexpected. By co-IP and Carm1-reporter assays, we determine that MPP7’s L27 domain is brought to the proximity of TAZ and YY1 through AMOT, and it strengthens the interactions between TAZ and YY1 to enhance *Carm1* transcription via TEAD and YY1 binding sites. The finding that TAZ, L27-TAZ, and YY1 can only activate the Carm1-reporter with both TEAD and YY1 binding sites intact and that no synergy was observed by their exogenous co-expression (though a higher level of reporter activity was found than each single expression) suggest that the primary mode of L27 is to enhance TAZ-YY1 interaction for higher occupancy efficiency on Carm1 promoter through DNA binding, instead of enhancing cooperativity of the transcription activating domains of TAZ and YY1.

### YY1 coordinates with YAP/TAZ to activate high levels of *Carm1* activation

YY1 contains distinct activation and repression domains and can function either way in a context-dependent manner (reviewed in Verheul et al., 2020). YAP/TAZ have also been shown to act as co-repressors via their DNA-binding partner TEAD (Kim et al., 2015). YY1 and YAP together repress cell cycle inhibitor genes in cancer cells by recruiting the repressive chromatin remodeler, EZH2 (Hoxha et al., 2020). In the context of MuSCs, YY1 has been studied as a repressor to regulate mitochondria function and glycolysis during MuSC activation (Chen et al., 2019). We were led to connect YAP/TAZ and YY1 by the co-existence of TEAD and YY1 binding sites in their common DEGs. Although we focused on the cooperative activation by YAP/TAZ and YY1 herein, they may cooperatively repress certain common DEGs. For example, 2 mitochondrial genes (*Slc25a29* and *Cmc1*; Table S1) are upregulated in the *Yy1* cKO and the *YazTaz* cKO and may be co-repressed by YY1 and YAP/TAZ. We chose the downregulated *Carm1* for an in-depth study because of YAZ/TAZ’s activator function in MuSCs (Judson et al., 2012; Sun et al., 2017; Tremblay et al., 2014) and *Carm1’s* role in MuSC renewal (Chang et al., 2018; Kawabe et al., 2012). Given the co-IP and Carm1-reporter results, the rescue of *Mpp7* cKO MuSCs by L27-TAZ but not TAZ suggests that the L27 domain of MPP7 is needed for TAZ (YAP) to cooperate with endogenous YY1 for MuSC renewal.

The logic of a multi-tiered control of *Carm1* expression, in addition to its post-transcriptional regulation (Chang et al., 2018), may lie in a checkpoint by the convergence of YY1’s and YAP/TAZ’s transcriptional programs. YY1 largely controls mitochondrial and glycolytic genes (Chen et al., 2019), whereas YAP/TAZ largely controls cell growth and proliferation genes (Sun et al., 2017 and herein). These cellular functions have to be coordinated for robust stem cell activation and progenitor expansion. Once these cellular conditions are coordinated, the convergence to renewal gene(s), e.g. *Carm1*, is set in motion. The inclusion of AMOT and MPP7 renders a mechano-checkpoint for MuSCs to sense their local regenerative physical environment and adjust the renewal rate accordingly. Lastly, both YY1 and YAP/TAZ pathways have also been extensively studied in cancer cells. Whether *Mpp7* and *Amot* contribute to cancers originating from MuSCs, e.g., rhabdomyosarcoma, deserves attention.

## Supporting information

Table S1 DEGs and cross comparison

Table S2 Reagents resource

---

**Figure S1. Additional data to support** Figure 1.

(A) Scheme for FACS isolation of YFP-marked Con and *Mpp7* cKO MuSCs from hindlimb muscles; FACS profiles to the right.

(B) Percentages of MPP7^+^ MuSCs from Con and *Mpp7* cKO immediately after FACS isolation.

(C) Representative IF images of PAX7 and LAMININ for Con and *Mpp7* cKO TA muscles 30 d (days) after TMX regimen without injury; white arrows, PAX7^+^ MuSCs. Quantification of PAX7^+^ SCs is to the right. N = 3 mice, each.

(D, E) Representative images of H&E histology (D) and IF (E) at 21 dpi. Black arrows indicate regenerated myofibers, and dashed lines, injury boundary in (D); white arrows, PAX7^+^ cells in (E); associated with Figures 1G and 1H.

(F) Representative images of EdU Click-reaction with IF of YFP; yellow arrowheads, EdU^+^YFP^+^ cells; white arrows, EdU^-^YFP^+^ cells; associated with Figure 1I.

(G) Representative IF images of PAX7 and MYOD with DAPI from single myofiber cultures; asterisks, PAX7^+^ cells; white arrows, PAX7^+^MYOD^+^ cells; yellow arrows, MYOD^+^ cells; associated with Figure 1J.

(H-J) Regimen to determine EdU incorporation and programmed cell death of MuSCs in culture in (H). (I, J) Percentages of EdU^+^ and cleaved-Caspase 3^+^ cells, respectively. DOXO was used to demonstrate cleaved-Caspase 3 reactivity.

Data information: Scale bars = 25 µm in (C-F, J). Error bars represent means ± SD. Student’s *t*-test (two-sided). n.s, *P* > 0.05; ***, *P* < 0.001.

**Figure S2. Additional data to support** Figure 2.

(A) IF of M-Cadherin (MCAD), N-Cadherin (NCAD), β-Catenin (CTNNB1), and PAR3 in Con and *Mpp7* cKO MuSCs immediately after single myofiber isolation; yellow arrowheads, apical side. Quantified fluorescent signals (AU) are to the right; total 30 cells in each group. N = 2 Con mice; N = 3 *Mpp7* cKO mice.

(B) Distribution pattern of V5-MPP7 variants in transfected *Mpp7* cKO MuSCs on single myofibers was determined by IF of V5; yellow arrowheads, apical side. Relative fractions of nuclear versus non-nuclear MPP7 are to the right: WT, 52 cells; △L27, 55 cells; △PDZ, 54 cells; △△SH3GUK, 52 cells; associated with Figure 2B.

(C) IF of AMOT in Con and *Mpp7* cKO MuSCs on single myofibers; yellow arrowheads, apical side. Quantification of fluorescence intensity (AU) is to the right of representative images; total 30 cells. N = 2 Con mice; N = 3 *Mpp7* cKO mice; associated with Figure 2K.

(D) Summarized models: Top, MPP7 and AMOT interaction domains. Same color-coded domains of each protein as in Figure 2A and D, with only the relevant domains labeled; * denotes S175 of AMOT. Bottom: AMOT level depends on MPP7.

(E) Western blots used in Figure 2D; red rectangles outline the cropped images used; asterisks, non-specific bands.

Data information: Scale bars = 10 µm in (A-C). (A, C): Error bars represent means ± SD; Student’s *t*-test (two-sided). B, Chi-square test was performed. n.s, *P* > 0.05; ***, *P* < 0.001.

**Figure S3. Additional data to support** Figure 3.

(A) Representative IF images of CARM1 in *Mpp7* cKO and *Amot* cKO MuSCs on single myofiber culture at 48 h; yellow arrowheads, apical side; associated with Figure 3F.

(B) Relative activities of the Carm1-reporter with V5-Mpp5 or V5-Mpp7 expression constructs in 293T cells; (-), empty vector. N = 3.

(C) Relative activities of the Carm1-reporter when co-transfected with WT and deletion mutants of Mpp7 expression constructs (x-axis) in 293T cells; (-), empty vector. N = 3.

(D) Venn Diagrams compare DEGs from myoblasts transduced by stabilized forms of YAP (S127A) and TAZ (S89A) (Sun et al., 2017) versus DEGs in *Mpp7* cKO (left) and *Amot* cKO (right) MuSCs; overlapping genes are underlined; associated with Figures 3C and 3D.

(E) TEAD binding site analysis of DEGs of the *Mpp7* cKO and the *Amot* cKO based on integrated experimental data sets (Keenan et al., 2019) and consensus sequence prediction (Fishilevich et al., 2017); associated with Figures 3C and 3D.

Data information: Scale bar = 10 μm in (A). (B, C) Error bars represent means ± SD. One-way ANOVA with Tukey’s post hoc test was performed. (D) Hypergeometric test was performed. n.s, *P* > 0.05; ***, *P* < 0.001.

**Figure S4. Additional data to support** Figure 4.

(A) Top 10 GO-term enriched pathways ranked by gene counts for the *YapTaz* cKO; gene counts and *P*-values on the right.

(B) Venn diagram for overlapping DEGs in the *Mpp7* cKO, the *Amot* cKO, and the *YapTaz* cKO; associated with Figure 4B.

(C) Enrichment of TEAD and YY1 binding sites in promoter regions of DEGs common between the *YapTaz* cKO and the *Mpp7* cKO, between the *YapTaz* cKO and the *Amot* cKO, and among all 3 cKOs; sources for analyses at the bottom; associated with Figure 4B.

(D) Quantification of IF signals (AU) of TAZ, YAP, AMOT, and CARM1 of 48 h cultured Con, *Mpp7* cKO, and *Mpp7* cKO MuSCs treated with MG132 (added 24 h prior to assay); (-), DMSO (solvent for MG132) was added as mock-control in parallel; 200 MuSCs from 2-3 mice in each group; associated with Figures 4C and 4D.

(E) Representative IF images of V5 and CARM1 of 48 h cultured *Mpp7* cKO MuSCs when transfected with V5-Taz and V5-Taz S89A expression constructs; associated with Figure 4H.

(F) Western blots used in Figure 4E; red rectangles outline the cropped images used. Data information: Scale bar = 10 μm in (E). (D) Error bars represent means ± SD. One-way ANOVA with Tukey’s post hoc test was performed. n.s, *P* > 0.05; *, *P* < 0.05; ***, *P* < 0.001.

**Figure S5. Additional data to support** Figure 5.

(A) MPP7-TAZ interaction is enhanced by WT AMOT, but disrupted by ΔPDM AMOT using the co-IP assay in 293T cells. Expression constructs and tagged epitopes for detection are indicated; (-), empty vector; quantification of co-IPed V5-TAZ to the right. N = 3.

(B) Mutation in the AMOT-interacting WW-domain of TAZ abolishes MPP7-TAZ interaction. Expression constructs and tagged epitopes for detection are indicated; (-), empty vector. WWm, WW domain of TAZ with a point mutation that disrupts its interaction with AMOT. N = 2.

(C) MPP7’s L27 domain has two L27 motifs, L27N and L27C. Taz-fusion constructs with only L27N or L27C were made. Taz-fusion constructs with L38S or L95S a.a. substitution in the L27 domain were also made; (-), empty vector. They were tested for activity on the Carm1-reporter; expression constructs in the x-axis. N = 3.

(D) gfp (control), V5-L27-Taz, and V5-PDZ-Taz expressing constructs were transfected into cultured *Mpp7* cKO MuSCs. IF of V5 and CARM1 were performed (representative images on the left) and signals (AU) of CARM1 were quantified; 200 MuSCs in each group.

(E) Western blots used in Figure S5A; red rectangles outline the cropped images used. Red asterisks indicate non-specific bands.

(F) Western blots used in Figure 5A; red rectangles outline the cropped images used.

(G) Western blots used in Figure S5B; red rectangles outline the cropped images used; red asterisks, non-specific bands.

Data information: Scale bar = 10 µm in (D). Error bars represent means ± SD. One-way ANOVA with Tukey’s post hoc test was performed in (A, B). n.s, *P* > 0.05; *, *P* < 0.05; ***, *P* < 0.001.

**Figure S6: Additional data to support Figure 6.**

(A) Percentages of nuclear IF signals (versus total IF signals) of MPP7 and AMOT for data in Figure 6D; treatments follow those in Figure 6D and are in x-axis; 200 MuSCs in each group.

(B) An example for how nuclear MPP7 and AMOT immunostaining signals were quantified. FACS-isolated YFP-marked MuSCs were used. DAPI was used for co-staining to define the area of the nucleus (yellow dashed lines), and YFP was used to define the total area of a cell (white dashed lines). The percentage of nuclear signal is that of the signal overlapped with DAPI divided by total signals within YFP-marked area. Same images were shown in Figure 6C (DMSO control); associated with Figures 6C-E.

Data information: Scale bar = 10 μm in (B). (A) Error bars represent means ± SD. One-way ANOVA with Tukey’s post hoc test was performed. ***, *P* < 0.001.

**Figure S7. Additional data to support** Figure 7.

(A) Venn Diagrams of DEGs between the *Yy1* cKO and the *YapTaz* cKO, the *Yy1* cKO and the *Mpp7* cKO, and the *Yy1* cKO and the *Amot* cKO. The data set of *Yy1* cKO was performed at 36 h of culture, from Chen et al. (2019).

(B) WT, TEAD binding site mutated, and YY1 binding site mutated Carm1-reporters were tested for response to the V5-Mpp7 expression construct; (-), empty vector. N = 3.

(C) WT Carm1-reporter was tested for response to L27-Taz or L27-Taz with a mutated WW domain (WWm, no longer interacting with AMOT), with or without YY1 co-expression (indicated in the x-axis); (-), empty vector. N = 3.

(D) Western blots used in Figure 7E; red rectangles outline the cropped images used.

(E) Western blots used in Figure 7F; red rectangles outline the cropped images used. Data information: Error bars represent means ± SD. Student’s *t*-tests (two-sided) were performed in (B). One-way ANOVA with Tukey’s post hoc test was performed in (C). n.s., *P* > 0.05; ***, *P* < 0.001.

## MATERIALS AND METHODS

### Mice

Animal treatment and care followed NIH guidelines and the requirements of Carnegie Institution, and approved by Carnegie Institutional Animal Care and Use Committee. *Mpp7^flox^* mice was generated via contractual service with ALSTEM Inc., and available upon request. *Pax7^CreERT2^* mice (Lepper et al., 2009) were donated by Dr. C Lepper. *Amot^flox^* mice (Shimono and Behringer, 2003) was obtained from Dr. J Kissil. *Yap^flox^*, *Taz^flox^* (Reginensi et al., 2013), and *Rosa26^YFP^* (Srinivas et al., 2001) mice were obtained from The Jackson Laboratory. Mice were genotyped by PCR using tail DNA by allele-specific oligonucleotides (information available upon request). Appropriate mating schemes were performed to obtain control and experimental mice stated in text, figures and legends. Both male and female mice were used and included in data analysis unless specified otherwise.

### Animal procedures

Mice (3-6 month of age) were administered intraperitoneally for 5 consecutive days with 250 μL tamoxifen (10 mg/mL corn oil; Millipore Sigma), followed by 3 days of chase. For muscle injury, mice were anesthetized and 50 μL of 10 μM cardiotoxin (CTX; Millipore Sigma) in PBS was injected into tibialis anterior (TA) muscles using the BD insulin syringe (Becton Dickenson). For EdU (5-ethynyl-2’-deoxyuridine; Millipore Sigma) incorporation in vivo, 10 μL of EdU (0.5 mg/ml in PBS) per gram of weight was used per intraperitoneal injection. Time lines of experimental procedure and muscle sample harvest are detailed in figures and legends.

### Histology and Immunofluorescence (IF)

TA muscles were fixed in 4% PFA (Electron Microscopy Sciences) immediately after harvesting. They were processed through 10% sucrose/PBS, 20% sucrose/PBS, and FSC 22 frozen section media (Leica) before mounted onto a cork and flash-frozen in liquid nitrogen cooled isopentane (VWR). Frozen samples were stored in -80°C until sectioning by a cryostat (Leica CM3050 S). Sections of 10 µm thickness were collected on Superfrost plus slides (VWR), dried, and stored at -20°C for future use. For histology, Hematoxilin Gill’s II and Eosin (H&E) were used following instructions of the manufacturer (Surgiopath), and mounted in Permount (VWR). For IF, sections were permeabilized with 0.6% TritonX-100/PBS for 20 min, blocked in Mouse on Mouse (M.O.M; Vector) blocking reagent, and then in blocking buffer (10% normal goat serum (Gibco) or normal donkey serum (Sigma-Aldrich), 10% carbo-free blocking solution (Vector) in PBS). Harvested single myofibers and cultured cells (see below) were fixed and permeabilized the same way and blocked in blocking buffer without M.O.M. reagent. Tissues and cells were incubated with primary antibodies overnight at 4℃ and secondary antibodies for 1 h at room temperature. DAPI was used to detect nuclei. EdU incorporation was detected by Click-iT Alexa Fluor 647 Imaging kit (Thermo Fisher Scientific). Brightfield microscope (Nikon Eclipse E 800), fluorescence microscope (Nikon E800), and confocal microscope (Leica TCS SP5) were used for imaging.

### MuSC isolation by fluorescence activated cell sorter (FACS)

YFP-labeled MuSCs were isolated by FACS from control and cKO mice specified in text, figures, and legends. Briefly, hindlimb muscles were minced and digested with 0.2% collagenase (Worthington Biochemical) for 90 min followed by 0.2% dispase (Thermo Fisher Scientific) for 30 min in 37℃ shaking water bath. Triturated muscle suspension was filtered through a 40 µm cell strainer (Corning) and subjected to isolation by BD FACSAriaIII. For culture, mononuclear cells were seeded on Matrigel (Corning) coated plates in growth media (DMEM with 20% FBS, 5% horse serum, 1% pen-strep, 1% glutamax (above from Gibco), 0.1% chick embryo extract (MPbio) and 2 ng/mL FGF2; R&D systems) for specified time in text and legends, before fixation and analysis.

### Single myofiber isolation

Single myofibers were isolated from extensor digitorum longus (EDL) muscles as described (Li and Fan, 2017). Isolated MuSCs were fixed in 4% PFA immediately or cultured in DMEM with 10% horse serum and 0.5% chick embryo extract for specified time in text, figures, and legends. To preserve MuSC projections, modifications were made to the procedure in (Kann et al., 2022). Knee tendon was cut prior to ankle tendon to remove the EDL muscle. Tugging and pulling were avoided to prevent muscle stretching and loss of projections. EDL muscles were digested in 2.6 mg/mL collagenase in DMEM with Y-27632 (50 μM; Tocris Bioscience) for 55 min in 37℃ shaking water bath, and transferred to DMEM with Y-27632 (50 μM) for trituration to liberate individual myofibers. These single myofibers were immediately fixed in 4% PFA. After fixation, myofibers and their associated MuSCs were subjected IF and imaging analysis.

### RNA-seq and analyses

For RNA-seq, 3-month old female mice were used. YFP-labeled MuSCs were purified by FACS and processed for RNA extraction using Direct-zol RNA Miniprep Kit (Zymo Research). Total RNA was processed by ribosomal RNA depletion using the Ribo-Zero rRNA Removal Kit (Illumina) and sequencing library generated using the TruSeq RNA Library Prep Kit (Illumina) with omission of PolyA selection. Raw data from FastQ were processed using standard method (Pertea et al., 2016) and the reads were mapped to the mouse mm9 genome. Differentially expressed gene (DEG) analysis was performed with DESeq2 (Love et al., 2014) with default parameters. Transcription factors binding sites in gene promotors were identified by using ChEA3 (Keenan et al., 2019) and GeneHancer prediction (Fishilevich et al., 2017). To cross-compare our DEGs were with *Yap/Taz* over-expression and *Yy1 cKO*, we extracted DEGs from (Sun et al., 2017) and (Chen et al., 2019), respectively for analyses. Significance of the rate of enrichment was assessed using hypergeometric test, and *P*-values stipulated in figures.

### Plasmid transfection of single myofibers and 293T cells

For single myofiber transfection, myofibers isolated from Con, *Mpp7* cKO or *Amot* cKO were cultured for 12 hr and transfected with indicated expression plasmids (Table S2) using TransfeX reagent (ATCC). Myofibers were then cultured and harvested at indicated time; 1 µM 4-OH-TMX (Tocris Bioscience) was added to sustain knockout efficiency. For 293T cells, indicated plasmids were transfected using Lipofectamine 3000 (Thermo Fisher Scientific). After 24 hr, cells were lysed by RIPA buffer (50 mM Tris-HCl [pH 8.0], 150 mM NaCl, 1% NP-40, 0.5% deoxycholic acid, and 0.1% SDS) supplemented with complete protease inhibitor cocktail (Roche) and 1 mM PMSF (Millipore Sigma) processed for Western blot detection.

### Co-immunoprecipitation (co-IP) and Western blot

Cell lysates from transfected 293T were incubated with anti-FLAG M2 magnetic beads (Millipore Sigma) or anti-V5 agarose beads (Millipore Sigma) at 4°C for 4 h or overnight. Beads were then washed 3 times with NP-40 cell lysis buffer (50 mM Tris-HCl [pH 8.0], 150 mM NaCl, 1% NP-40 supplemented with 0.5 mM dithiothreitol) and 1 time with PBS. 5% input and immunoprecipitated fractions were boiled in Laemmli SDS sample buffer (Thermo). Protein samples were processed for SDS-PAGE (4-15% gel; Bio-Rad), transferred to PDMF membrane (Bio-Rad) for detection using rabbit anti-HA, rabbit anti-V5, or rabbit anti-FLAG antibodies (Cell Signaling), followed by HRP-conjugated goat anti-rabbit antibodies (Bio-Rad). ECL substrate (Thermo Fisher Scientific) was used for detection. Exposure and images were performed using LI-COR Fc imager (LI-COR biosciences).

### Luciferase assay

Carm1 promoter region (-630 to +15) was cloned to pGL4 luciferase reporter vector (Promega) to be the Carm1-reporter. Carm1 promoter region was analyzed by PROMO (Farre et al., 2003; Messeguer et al., 2002) and JASPAR (Castro-Mondragon et al., 2022) to identify TEAD and YY1 binding sites. Carm1-reporter with TEAD and YY1 binding site mutated were generated by PCR and the nucleotide sequences are indicated in figures. The Carm1-reporter or its mutated reporters was co-transfected into 293T with the pRL-TK plasmid (Promega) expressing renilla for normalization. Combinations of cDNA expression plasmids (Table S2) were indicated in figures and legends. Twenty-four h after transfection, 293T cells were harvested and luciferase and renilla activities were detected using the Dual-luciferase reporter assay kit (Promega) in a Glowmax 20/20 luminometer (Promega). The 8XGTIIC-luciferase vector (Dupont et al., 2011) was used as the TEAD-reporter using the same procedure. For reporter assays, 3 independent biological replicates were performed for each combination of reporters and cDNA expression plasmids.

### Pharmacological treatments

For EdU incorporation in cultured MuSCs, EdU was mixed in media to a final concentration of 10 µM for 24 hr, followed by the Click-reaction (Thermo Fisher Scientific) for detection. FACS-isolated MuSCs were immediately treated with DMSO (mock-treatment), Jasplakinolide (100 nM; Tocris Bioscience) or Narciclasine (100 nM; Tocris Bioscience) at for 2 hr, cytospun to coverslip, and processed for IF. For activated MuSCs, they were cultured for 48 h after FACS-isolation, and treated by DMSO, Blebbinstatin (10 µM; Tocris Bioscience), Cytochalasin B (10 µM; Tocris Bioscience), and Y-27632 (10 µM) for 2 hr and processed for IF.

### Quantifications and statistical analysis

For cryosections, ≥ 50 cells from 10 sections per animal (5 animals per genotype) were imaged using a Nikon E800 fluorescence microscope at 40X magnification. For MuSC fractions on single myofibers, total 150-200 cells were assessed from 2-3 animals using the same microscopy above. For quantification of IF signal intensity, 50 cells on myofibers (from 2-3 mice) or 200 dish-cultured MuSCs (from 2-3 mice) were imaged at 63x/1.4 oil fluorescent objectives on a Leica TCS SP5 confocal microscope. Gain and exposure settings were consistent between experiments. Z-stacks were collected to capture full objection lengths of MuSCs. Images were exported to Fiji and CellProfiler for analysis.

For quantification of IF signal intensity and nuclear vs. cytoplasmic distribution, single myofibers or cultured cells were IF-stained for protein of interest (i.e., MPP7, AMOT, YAP, TAZ, CARM1, or tagged epitopes), YFP and DAPI, and multi-channel images were acquired. A custom CellProfiller pipeline was used to set threshold on YFP and DAPI channels to identify primary and secondary objects, respectively. The primary objects from YFP channel were then used as masks on protein-of-interest channel images, and the integrated intensity of the masked image was used for total signal intensity. The secondary objects from DAPI channel were used as masks on protein-of-interest channel images and the intensity of the masked part was quantified for nuclear signal intensity.

For co-IP quantification, the samples were analyzed in three independent biological replicates. Intensity of blotting bands were measured using Fiji. Co-IPed target proteins were then normalized to their primary IP proteins.

For statistics, error bars represent means ± SD. Data analyses were performed by Prism 9 software. Data comparison of two independent groups was performed by two-tailed unpaired Student’s t test. Multiple group analysis was performed using one-way ANOVA followed by the Tukey or Dunnet post hoc test for multiple comparison per figure legends. To test significance of cell population fraction, the total SC population over all the experimental repeats were included and comparisons were performed by Chi-squared test. ***, *P* ≤ 0.001; **, *P* ≤ 0.01; *, *P* ≤ 0.05; n.s, not significant, *P* > 0.05. For RNA-seq, *P* < 0.05 was considered significant. To calculate the statistical significance of overlapped genes in Venn diagrams, *P*-values were calculated based on hypergeometric test.

## Data Availability

RNA-seq data in this study have been deposited to Gene Expression Omnibus (GEO) database under the accession GSE241340.

## Acknowledgment

We thank Dr. Lydia Li for the initial finding. Dr. Wanjin Hong for sharing the Taz (WT and WWm) plasmids. Dr. Frederick Tan for training in transcriptome analysis, Allison Pinder for assistance in RNA-seq, Dr. Mahmud Siddiqui for assistance in microscopy, Liangji Li and Haolong Zhu for assistance with FACS and single myofiber culture, and Colin Bylieu for tail PCR genotyping. We also thank the Carnegie rodent facility crew for taking care of mouse strains. C.-M.F. is supported by the NIH (R01AR060042, R01AR071976, and R01AR072644) and the Carnegie Institution for Science. A.S. is supported by the NIH (R01AR072644).

## Author Contributions

A.S. and C.-M.F. conceived and designed the study and wrote the manuscript. A.S. carried out all the experiments and data analyses. J.L.K. provided comments and suggestions, crucial expression plasmids, the *Amot*^flox^ mouse, and extensive manuscript editing.

## Notes

The authors declare no conflict

https://www.ncbi.nlm.nih.gov/geo/query/acc.cgi?acc=GSE241340

### Competing Interest Statement

The authors have declared no competing interest.

