## Supplementary figures and images for "The L27 Domain of MPP7 enhances TAZ-YY1 Cooperation to Renew Muscle Stem Cells"

### supplemental figures

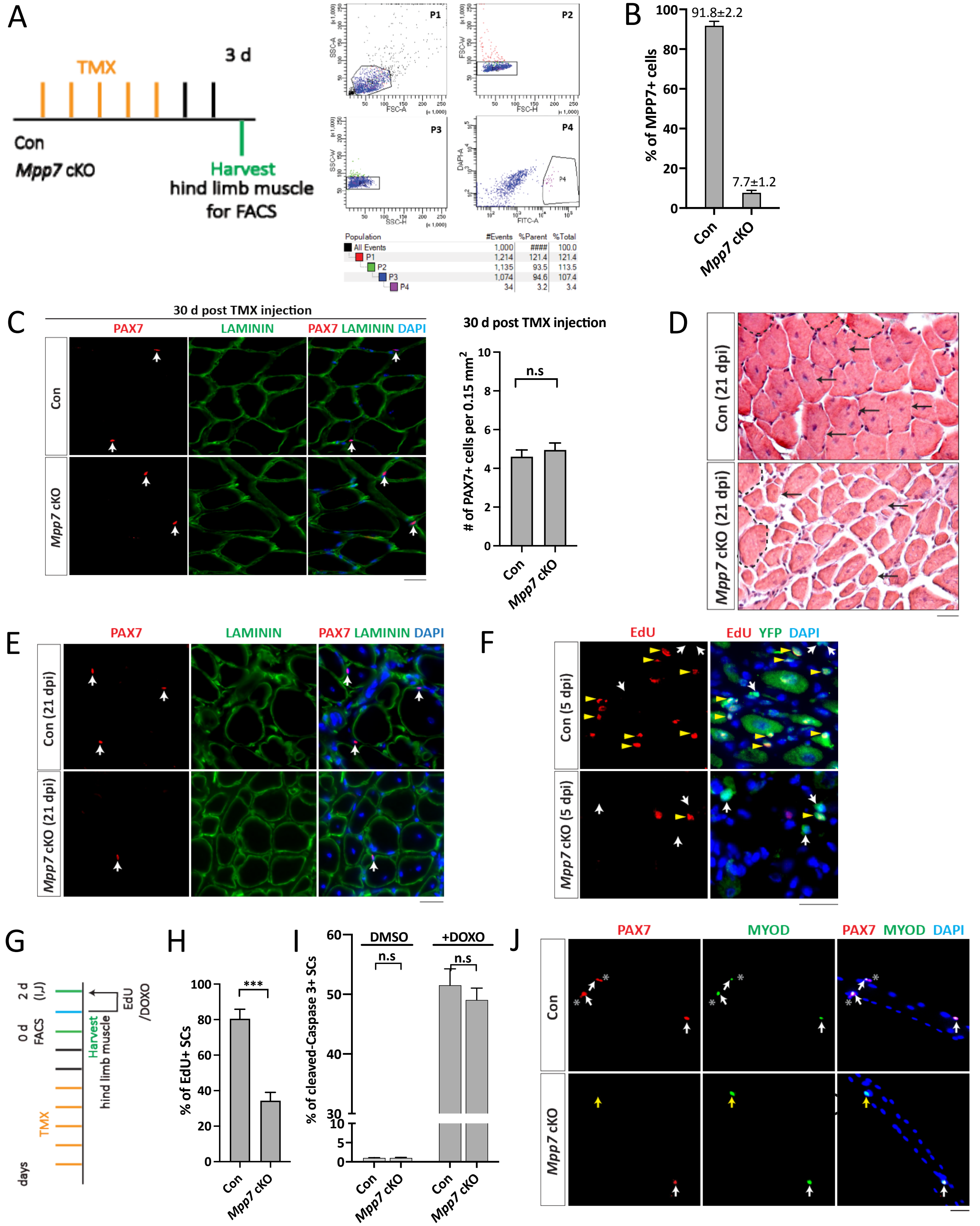

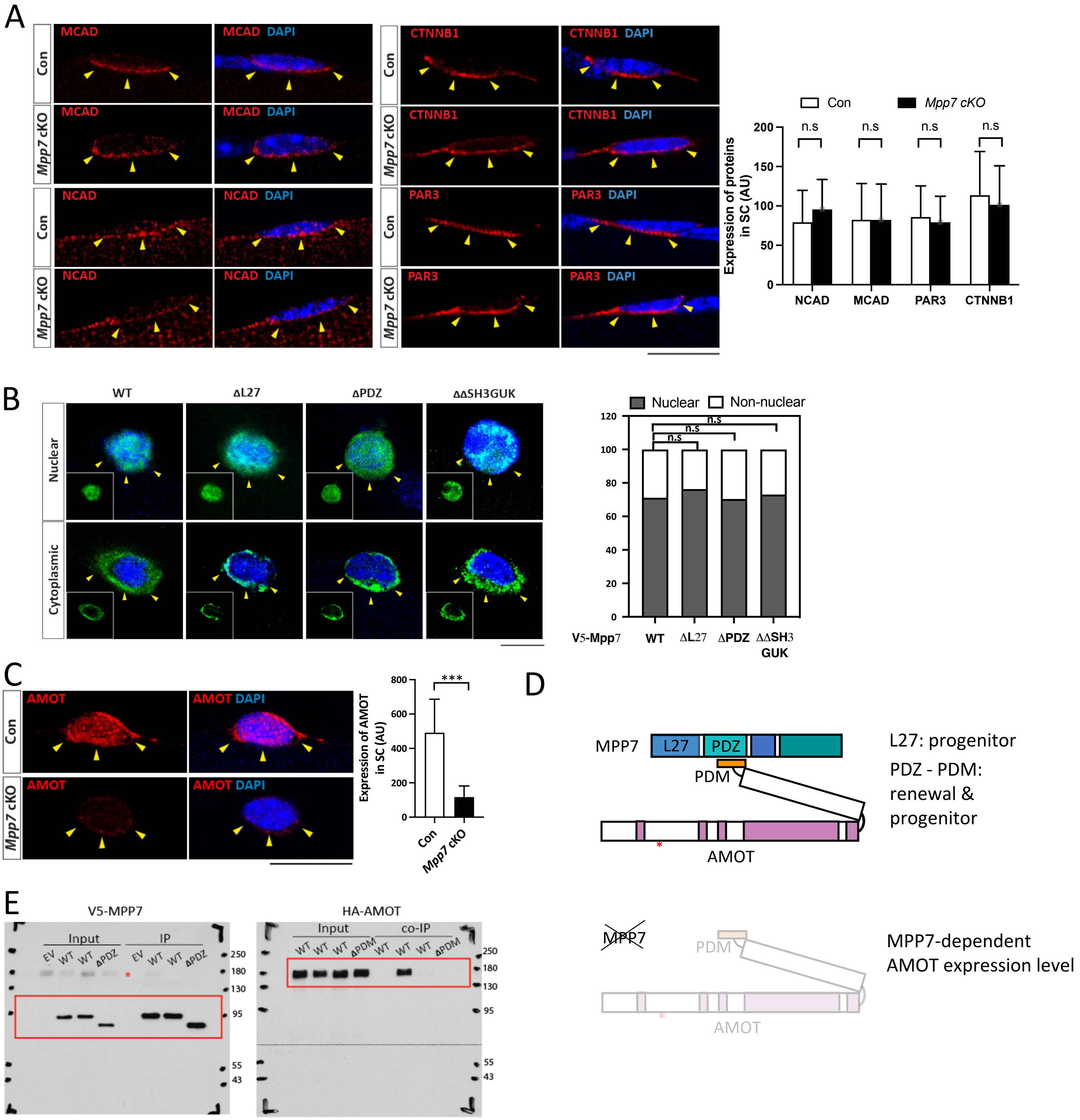

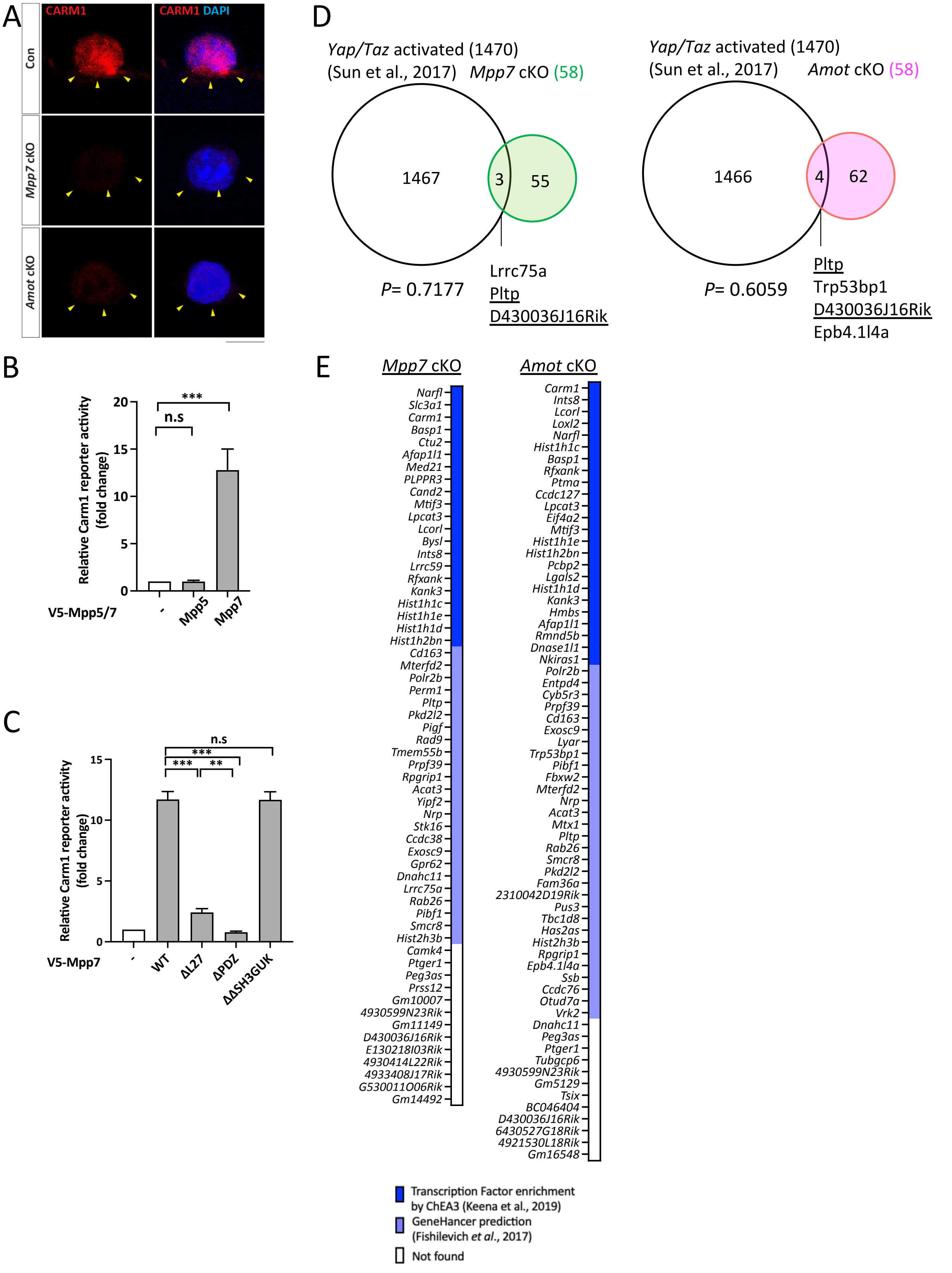

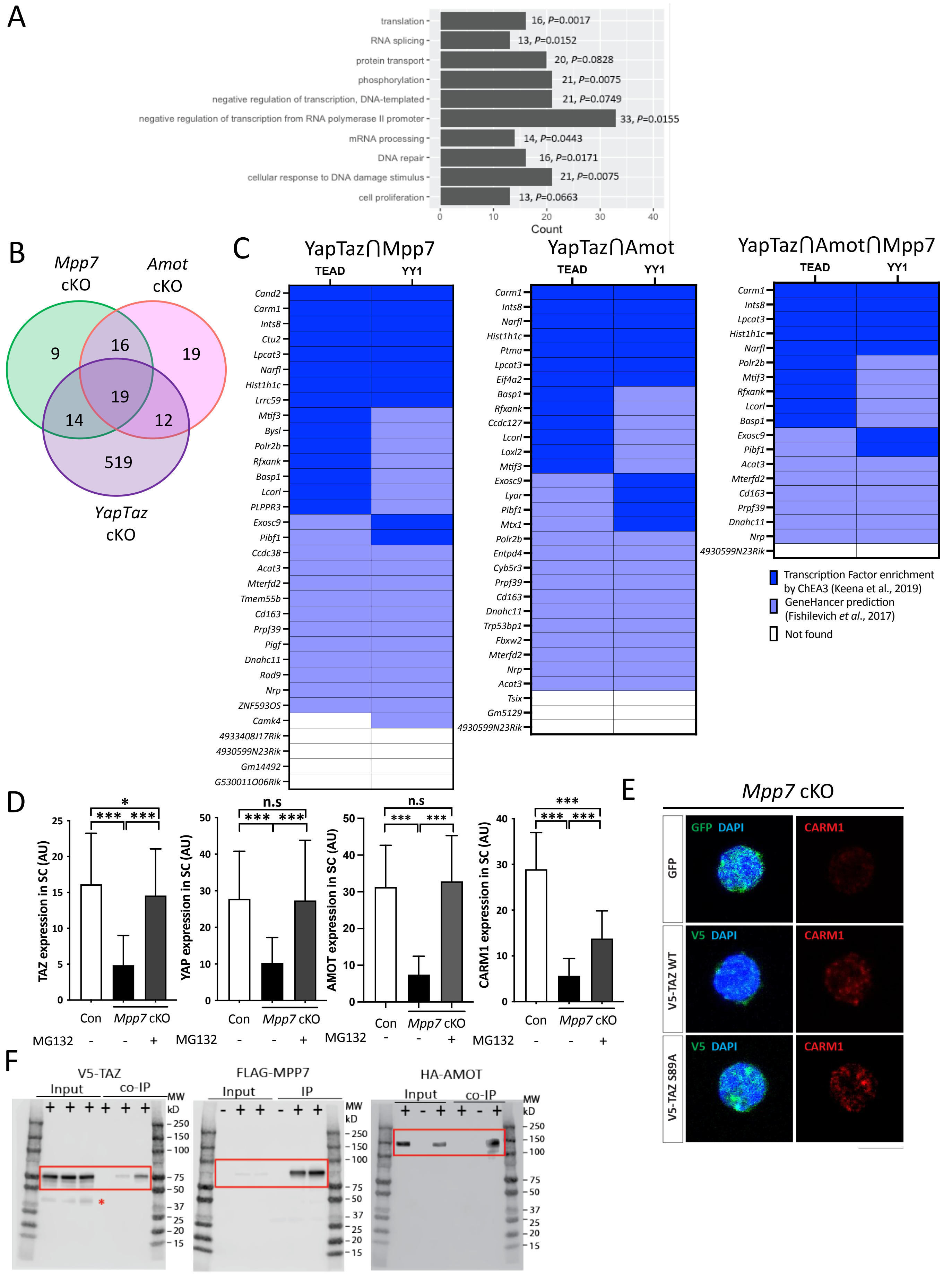

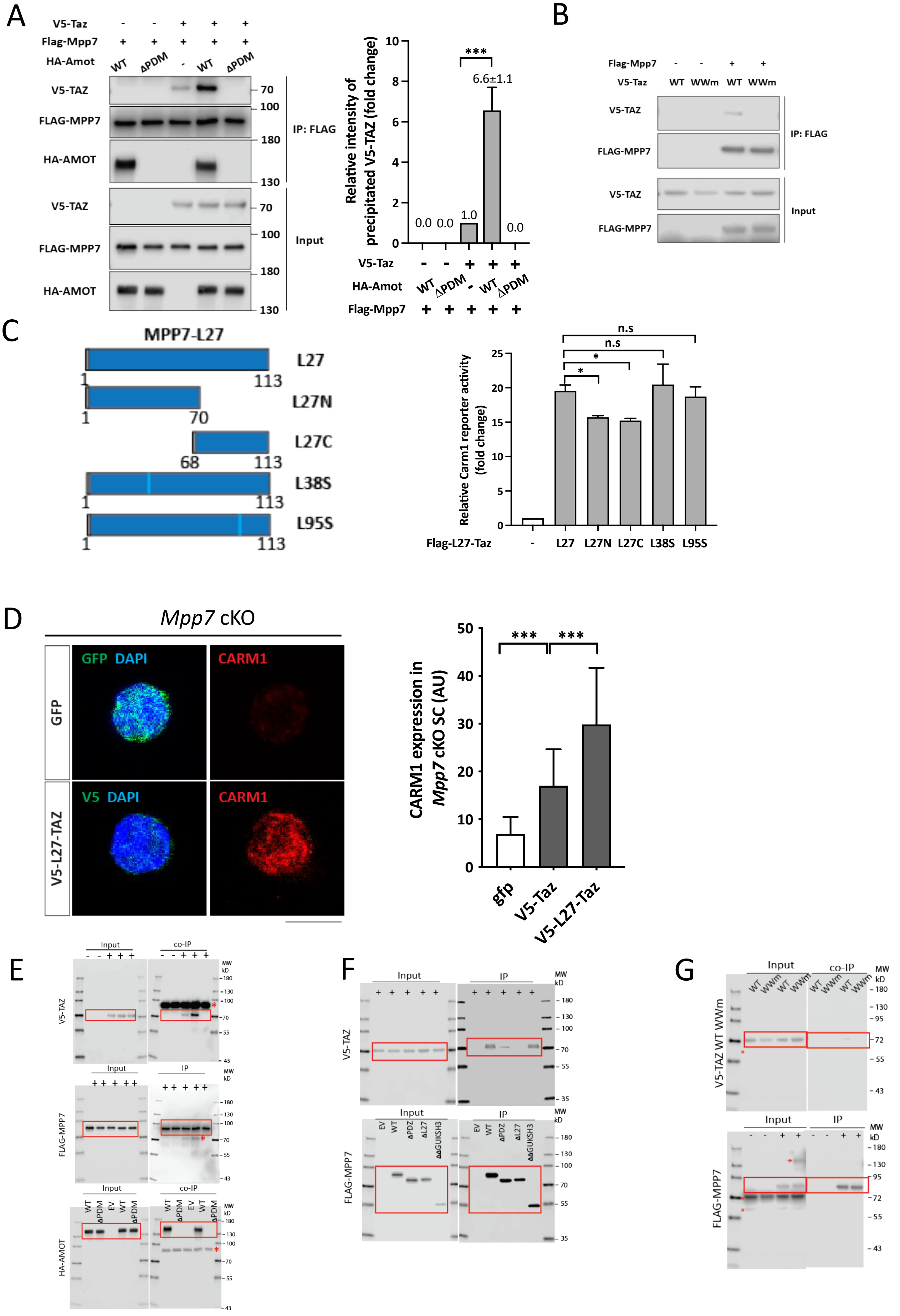

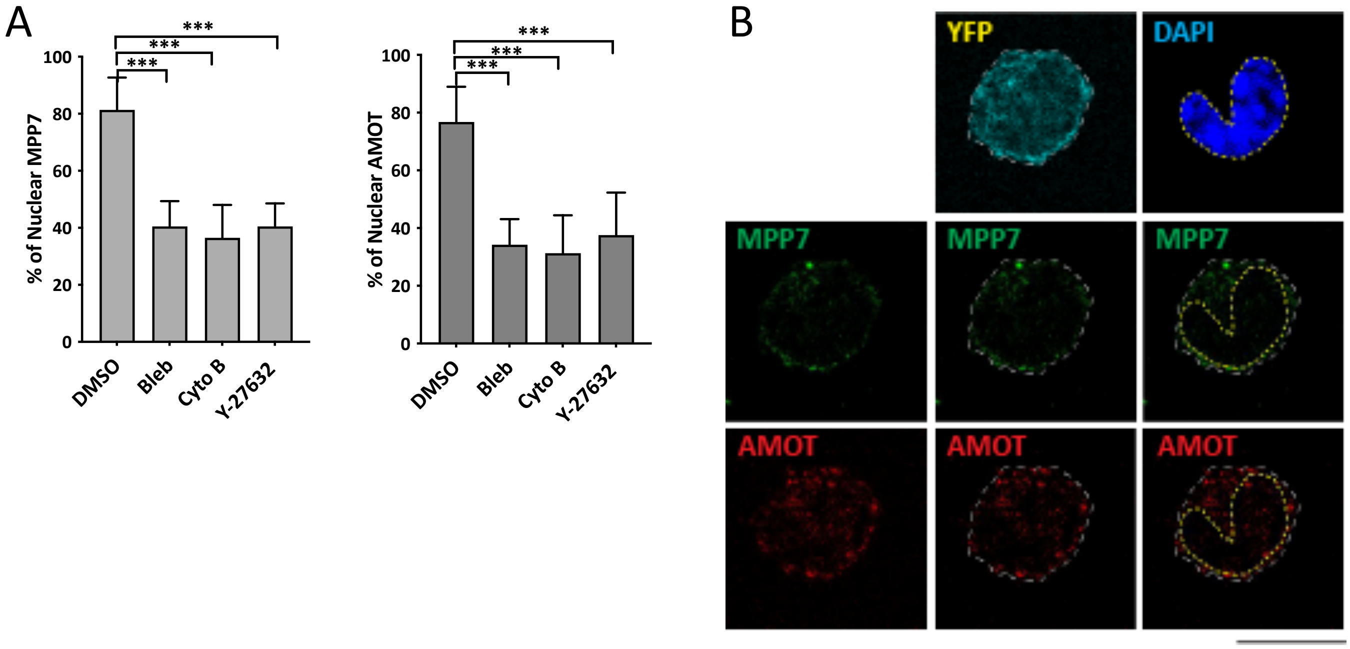

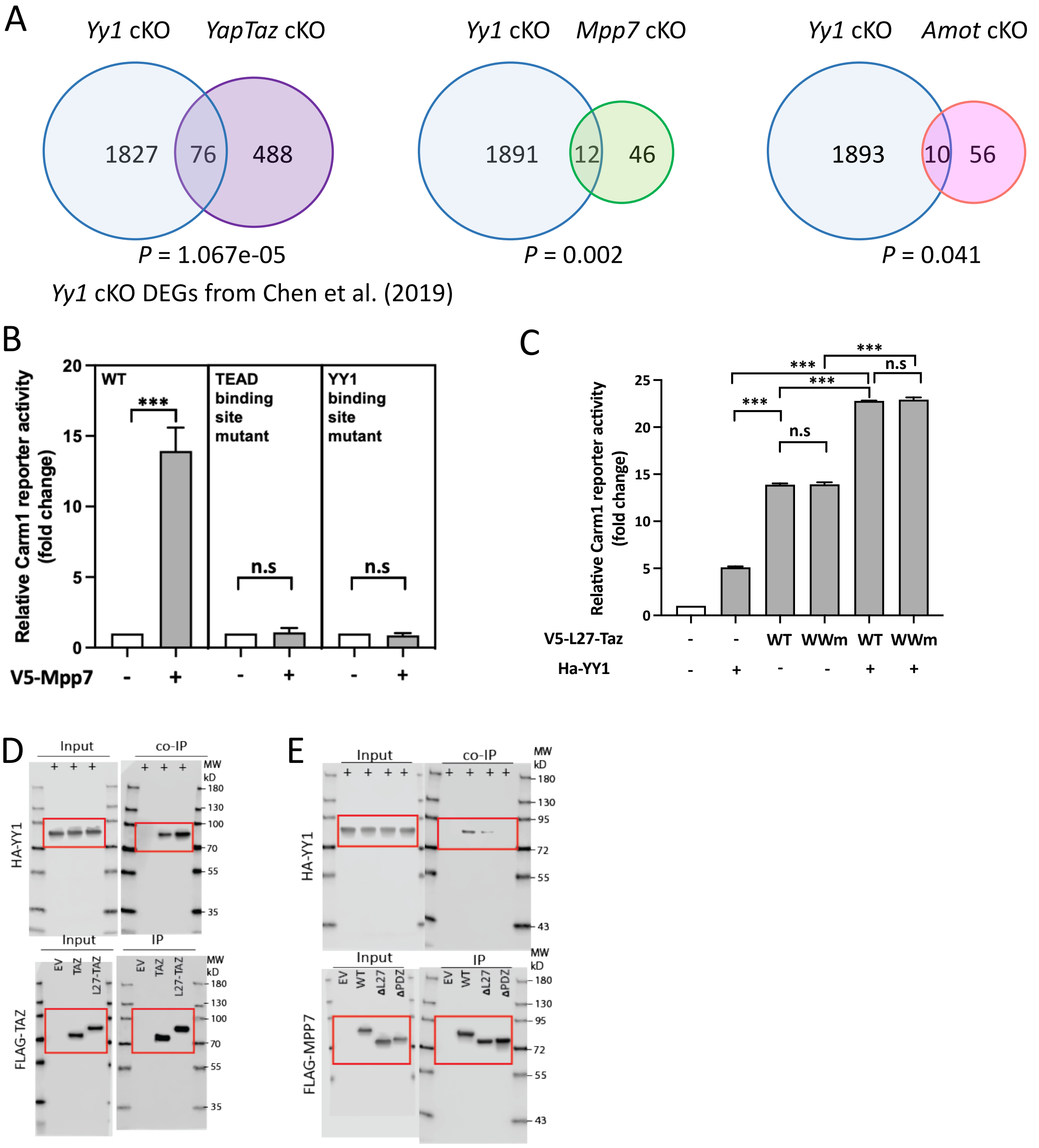
